# Comparing fine-scale mutation and recombination landscapes in rhesus macaque (*Macaca mulatta*) populations of Chinese and Indian descent inferred from both short- and long-read sequencing data

**DOI:** 10.64898/2026.05.26.727910

**Authors:** Gabriella J. Spatola, Cyril J. Versoza, Vivak Soni, Erangi J. Heenkenda, Jeffrey D. Jensen, Susanne P. Pfeifer

## Abstract

Genomic diversity amongst primates is fundamentally shaped by species- and population-specific rates of mutation and recombination. In this study, we infer fine-scale mutation and recombination rate maps for the rhesus macaque (*Macaca mulatta*) — the most widely used non-human primate model in biomedical research — leveraging both short-(Illumina) and long-(PacBio HiFi) read sequencing data from two distinct populations of Chinese and Indian descent. Thereby, we draw comparisons between the rates estimated from each dataset, highlighting both biologically meaningful variation between these populations as well as artefactual discrepancies likely arising from systematic biases and differences in the utilized sequencing technologies. Consistent with previous observations in humans, broad-scale features of the recombination landscape are well-conserved between the two populations, but significant differences exist at the finer scales. Notably, we find evidence for a high rate of turnover in recombination hotspots over a short evolutionary time span, resulting in population-specific recombination maps in which the vast majority of the >30,000 identified recombination hotspots in one population are inactive in the other population. Given that mutation and recombination rates are necessary components for the interpretation of other diversity-shaping processes and events, including those characterizing both the underlying demographic and selective histories, the incorporation of these population-specific maps into future models will improve our understanding of the evolutionary genomics of the species. Additionally, these maps will serve as a fundamental component of future genome-wide association and fine-mapping studies of disease traits in this biomedical model system.

## Introduction

Spontaneous mutation and meiotic recombination in the germline are two fundamental evolutionary processes, responsible for creating and shuffling genomic variation. Mutations that newly arise in an individual, known as *de novo* mutations, originate from DNA damage or replication errors that occurred in parental germ cells or during early embryonic development that were not properly repaired (Crow 2000). Recombination generates novel combinations of mutations via homology-directed repair of targeted DNA breaks, with repair outcomes manifesting as crossovers, where homologous chromosome segments are exchanged, or non-crossovers (also referred to as gene conversion), where short tracts of sequence are non-reciprocally transferred between homologs (Johnston 2024). In primates, targeted DNA breaks are primarily regulated by the sequence-specific binding of the zinc-finger protein PRDM9, leading to areas of the genome with significantly increased recombination, i.e., hotspots (Baudat et al. 2010; Myers et al. 2010; Parvanov et al. 2010), the location of which appears to evolve rapidly in the primate lineage (Auton et al. 2012; Schwartz et al. 2014; Stevison et al. 2016; Pfeifer 2020a).

Recombination interacts with other evolutionary processes in important ways; for example, by reshuffling genetic variation, beneficial mutations may be consolidated on to a common genetic background, or linkage between beneficial and deleterious alleles may be disrupted, thereby facilitating natural selection (Muller 1964; Hill and Robertson 1966; Felsenstein 1974). Moreover, the local rate of recombination impacts the extent of genetic hitchhiking — most notably background selection — affecting the genome (Maynard Smith and Haigh 1974; Charlesworth et al. 1993; and see the review of Charlesworth and Jensen 2021). Variation in recombination and mutation rates also plays a key role in accurately reconstructing population history, particularly in dating the timing of inferred events, and a neglect of this heterogeneity can lead to both mis-inference and increased uncertainties in parameter estimates (Nicolaisen and Desai 2013; Johri et al. 2022b; Soni et al. 2024). Additionally, as the inference of natural selection is highly dependent on the accurate modeling of demography, recombination and mutation rate analyses are crucial first steps towards any general understanding of a given species from a genomic perspective (Johri et al. 2022a; Jensen 2023).

Rhesus macaques (*Macaca mulatta*) are among the most well-studied primates in evolutionary and biomedical research as a result of their widespread geographic territory (ranging from Afghanistan, Pakistan, and India on the western edge of their range to the Chinese coastline in the east; Groves 2001), relatively close genetic relationship and strong physiological similarities to humans, and suitability for captive breeding programs (see the review by Cooper et al. 2022). Indeed, rhesus macaques maintained at NIH-funded primate research centers play a central role in biomedical studies as translational models (for aging, behavior, cancer, cardiovascular disease, endocrinology, development, neurobiology, and metabolism), as well as for the study of infectious diseases of various etiologies (including Ebola, HIV/SIV, herpes viruses, influenza, SARS-CoV2, tuberculosis, and Zika) and for general vaccine development (see Rogers 2022 and references therein). As might be expected given the extensive habitat range of the species, considerable subdivision has been observed between populations, with particularly high genetic differentiation between individuals of Chinese and Indian descent (Hernandez et al. 2007; Heenkenda et al. 2026; Maruki et al. 2026). These populations differ in several important aspects of their phenotypes, most notably with regards to their size (Clarke and O’Neil 1999), behavior (Champoux et al. 1997), genetic composition at major histocompatibility loci (Doxiadis et al. 2001, 2003; Viray et al. 2001), as well as their susceptibility to diseases (Trichel et al. 2002) — all of which can impact the outcomes of biomedical studies. Research on rhesus macaques in the United States has historically been focused upon individuals of both Chinese and Indian descent; however, importation of individuals from India ended in the late 1970s and, while the majority of research to date remains focused on individuals of Indian descent, individuals since introduced into the research colonies are primarily of Chinese origin (Rogers 2022).

Prior studies of the rates and patterns of germline mutation in the species have largely focused on tracking the occurrence of *de novo* mutations within pedigrees (Wang et al. 2020; Bergeron et al. 2021). These studies have provided important insights into the average point mutation rate across the genome (0.58 × 10^-8^ – 0.77 × 10^-8^ per base pair per generation [/bp/gen]) necessary for the scaling of phylogenetic events. Yet, knowledge of the fine-scale mutation rate heterogeneity required to accurately infer demographic histories and construct population-specific evolutionary baseline models remains elusive. Similarly, direct estimates of recombination rates exist for the species, tracing crossover events through extended pedigrees (Rogers et al. 2006; Versoza et al. 2024). The first such coarse-scale pedigree-based genetic linkage map was built from ∼250 human microsatellites, with an average marker spacing of 9.3 cM (Rogers et al. 2006). More recently, a genetic linkage map was released based on whole-genome data from four three-generation pedigrees, offering a median resolution of 22.3 kb (Versoza et al. 2024). Both maps covered between 2,048 and 2,357 Mb of the autosomal genome, with average point estimates of 0.82–0.88 cM/Mb in colony-born Indian rhesus macaques. In contrast to mutation, indirect fine-scale estimates of recombination rate were inferred for rhesus macaques from patterns of linkage disequilibrium (LD); however, these estimates vary markedly based on population, approach, and scale. For example, inferring rates of recombination between markers genotyped in a captive population of Indian descent, Xue et al. (2016, 2020) reported fine-scale estimates ranging from 0.433 ± 0.333 cM/Mb to 0.448 ± 0.286 cM/Mb, whereas Terbot et al. (2025a) estimated a much higher genome-wide fine-scale recombination rate of 0.79 cM/Mb in wild Chinese rhesus macaques. As LD-based maps capture long-term, historically-averaged recombination patterns in a population, they may be influenced by additional evolutionary processes, including the population-specific demographic history. Crucially, while Terbot et al. (2025a) explicitly accounted for this factor by incorporating a well-fit demographic history of the Chinese population in their study, Xue et al. (2016, 2020) neglected to take the history of the Indian population into account. Moreover, in contrast to Terbot et al. (2025a) who made their map for the Chinese population publicly available to the community, Xue et al. (2016, 2020) failed to do so for their studied Indian population. Consequently, the extent to which the discrepancy in rate estimates is driven by differences in approach vs. biologically meaningful population-specific changes to the recombination landscape — as have, for example, been observed for human populations (e.g., Hinch et al. 2011; Spence and Song 2019) — remains unknown.

As recombination and mutation rates vary across the genome (Stapley et al. 2017), the genomic resolutions and quality of the data being analyzed may also affect study results. Notably, the majority of studies investigating fine-scale mutation and recombination rates were based on short-read data, with Illumina being the most common sequencing technology applied, providing researchers with billions of short (∼100–150 bp) reads at an affordable cost (Slatko et al. 2019). However, ongoing scientific progress has resulted in the recent emergence of high-fidelity long-read sequencing, producing highly accurate (>99.9%) reads with lengths of up to 15–20 kb, thus offering both an unprecedented genomic resolution (including highly repetitive and GC-rich regions difficult to study with short-read data) as well as a greater accuracy of resolved haplotypes (Amarasinghe et al. 2020; Browne et al. 2020; De Coster et al. 2021; Espinosa et al. 2024). This in turn provides considerable advantages, particularly for recombination rate inference; yet, to the best of our knowledge, no previous studies have applied long-read sequencing to investigate recombination in an LD-based framework.

Here, we characterize population-specific differences in the mutation and recombination landscapes of the two rhesus macaque populations most widely used in research, inferred from newly generated PacBio HiFi long-read data from 20 individuals of Chinese and Indian descent. We further compare our observations with results obtained from publicly available Illumina short-read data to gain a better understanding of how the application of short-versus long-read sequencing technologies might lead to systematic differences in mutation and recombination rate inference. Given the importance of fine-scale mutation and recombination rate maps for the interpretation of association and linkage studies targeting traits of biomedical relevance, the maps generated from this uniquely high-quality data will thus aid future research in this important non-human primate model for human health and disease.

## Materials and Methods

### Animal subjects

Rhesus macaques were housed in indoor or outdoor social housing at the Oregon National Primate Research Center (ONPRC). All husbandry practices conducted are performed in accordance with federal guidelines and regulations as stated in the National Institutes of Health Guide for the Care of and Use of Laboratory Animals. ONPRC is accredited by the Association for Assessment and Accreditation of Laboratory Animal Care, International. Buffy coat samples were previously collected and stored under Oregon Health and Science University (OHSU) IACUC protocol #IP00000367.

### Samples and sequencing

We extracted DNA from previously collected buffy coat samples of a cohort of 20 unrelated rhesus macaques, comprising ten individuals of Chinese descent and ten individuals of Indian descent, and sheared each DNA sample to a target fragment size of ∼10–20 kb using a Megaruptor 3 (Diagenode, Liège, Belgium). Afterward, we purified the fragmented DNA with SMRTbell cleanup beads and constructed sequencing libraries with the SMRTbell Prep Kit 3.0. We performed size selection on a Pippin HT system (Sage Science, Beverly, MA, USA), enriching for fragments with lengths between 10 and 25 kb, and evaluated fragment size distributions on a Femto Pulse system (Agilent, Santa Clara, CA, USA). We then used the PacBio Sequel II Sequencing Kit 3.1 to bind sequencing polymerase to the SMRT libraries for HiFi sequencing. To ensure optimal SMRT cell loading, we quantified each library with a Qubit HS assay (Invitrogen, Carlsbad, CA, USA) prior to loading the libraries onto Revio SMRT Cells. Lastly, we performed HiFi sequencing in CCS mode with 24-hour movie collections.

### Long-read data

We processed the data from each sample by first converting the reads into .fastq format using the *bam2fastq* function in pbtk v.3.4.0 (https://github.com/PacificBiosciences/pbtk) and then quality-controlling the reads using fastplong v.0.2.0 (https://github.com/OpenGene/fastplong). In brief, we checked for remnant adapters and low-quality sequences (using the “*-u* 40 *-q* 20” flags) but none were detected; however, a total of 502 reads (0.001%) were removed across all samples as they were < 1 kb (using the “*-l* 1000” flag). We used minimap2 v.2.26 (Li 2018) with the “*-ax* map-hifi” preset to align the remaining long-reads from each sample to the rhesus macaque genome assembly, rheMac10 (GenBank assembly: GCA_003339765.3; Warren et al. 2020). Using NVIDIA Parabricks v.4.4.0-1 (O’Connell et al. 2023), we identified single nucleotide polymorphisms (SNPs) in each sample with DeepVariant v.1.6.1 in the “pacbio” mode (Poplin et al. 2018). Afterward, we jointly genotyped samples using GLnexus v.1.4.1 (Yun et al. 2020), limiting the dataset to those sites genotyped in all samples. To reconstruct haplotypes, we phased the dataset using WhatsHap v.2.3 (Martin et al. 2016). Lastly, to determine which regions of the genome were accessible to our study, we first calculated the depth of coverage at each position for each sample using the *genomecov* function implemented in BEDTools v.2.30.0 (Quinlan and Hall 2010) and then created an accessibility mask, requiring a coverage of ≥ 2 for each sample.

### Short-read data

To obtain additional information about polymorphisms segregating in the two populations, we downloaded genotype information from 888 individuals available in the mGAP database (Bimber et al. 2019) whose whole genomes have been sequenced to high coverage. We first limited this dataset to high-quality biallelic SNPs by setting genotypes with a quality score (*GQ*) < 20 to missing and excluding any sites at which > 50% of the individuals exhibited missing genotypes from the dataset using BCFtools v1.14 (Danecek et al. 2021). We then estimated pairwise kinship coefficients using KING v.2.3.2 (Manichaikul et al. 2010) and removed one individual from each closely related pair using PLINK2 (Chang et al. 2015), based on a threshold (*--king-cutoff*) of 0.177 corresponding to first-degree relatedness (Figure S1). Afterward, we re-calculated minor allele counts and excluded any sites with minor allele counts < 5. This produced a final population-scale dataset of 28.8 million SNPs segregating with high-confidence in 658 rhesus macaques of Chinese and Indian descent. This dataset was used in the inference of neutral divergence and mutation (see the section entitled “Inferring fine-scale rates of neutral divergence and mutation”).

To have a matched dataset of short-read whole-genome sequences from rhesus macaques for comparison with our long-read dataset used in the inference of recombination, we randomly selected 10 unrelated individuals each from the Chinese and Indian populations and downloaded publicly available data for these individuals using the Sequencing Read Archive (SRA) Toolkit’s *prefetch* and *fasterq-dump* modules v.3.2.1 (https://github.com/ncbi/sra-tools) together with their accession numbers (Chinese individuals: SRR1944106, SRR1964571, SRR1964575, SRR1964578, SRR5626558, SRR5626567, SRR5626569, SRR5626570, SRR5626571, SRR5626573; Indian individuals: SRR1927125, SRR1929278, SRR1929310, SRR1929386, SRR1944126, SRR5628082, SRR5628096, SRR5628144, SRR5628181, SRR7865702). These individuals were previously sequenced to an average coverage of ∼30× (range: 24.3·–39.7·). We mapped the sequence reads from each individual to the rhesus macaque genome assembly, rheMac10 (GenBank assembly: GCA_003339765.3; Warren et al. 2020), using the NVIDIA Parabricks v.4.4.0-1 (O’Connell et al. 2023) wrapper for BWA-MEM (Li 2013) *fq2bam*. We then called, jointly genotyped, and filtered variants following the recommendations for variant discovery by the Genome Analysis Toolkit (GATK) v.4.2.6.1 (van der Auwera and O’Connor 2020). Specifically, we called sites in each individual using GATK’s *HaplotypeCaller*, setting (a) the “*--pcr-indel-model*” parameter to CONSERVATIVE as the library preparation protocol may have used PCR, (b) the “*--heterozygosity*” flag to 0.00328 and 0.00209 for the Chinese and Indian individuals, respectively, based on previously reported levels of genetic diversity in the two populations (Xue et al. 2016), (c) the “*-ERC*” parameter to BP_RESOLUTION to obtain genotype information for both variant (polymorphic) and invariant (monomorphic) sites, and (d) the “*--minimum-mapping-quality*” parameter to 40 to limit calling to regions with high-quality mappings. We then combined individual call sets using GATK’s *CombineGVCFs* and jointly genotyped all individuals using GATK’s *GenotypeGVCFs* (with the “*-all-sites*” flag enabled). Afterward, we used GATK’s *SelectVariants* to limit the dataset to fully genotyped (“*AN* == 20”), biallelic (“*--restrict-alleles-to* BIALLELIC”) SNPs (“*--select-type-to-include* SNP”) in each population and removed any sites fixed for the alternate allele (i.e., those with a minor allele frequency of 1) using VCFtools v.0.1.14 (Danecek et al. 2011) with the “*--maf* 0.001 *--max-maf* 0.999” parameters. Next, we applied the “hard filter” criteria recommended for non-model organisms without any curated “gold standard” variant call set that could be used to train GATK’s machine-learning algorithm. To this end, we used GATK’s *VariantFiltration* tool, excluding any sites where the quality by depth (*QD*) was < 2.0, the Phred-scaled confidence score that a variant exists at the site (QUAL) was < 30.0, the Phred-scaled *P*-value using Fisher’s exact test (*FS*) to detect strand bias was > 60.0, the symmetric odds ratio (*SOR*) to detect strand bias was > 3.0, the mapping quality (MQ) was < 40.0, or the *Z*-scores from the Wilcoxon rank sum tests of position bias (*ReadPosRankSum*) and alternative versus reference read mapping qualities (*MQRankSum*) were < –8.0 and < –12.5, respectively. Following the guidelines described in previous recombination rate estimation studies in primates (Auton et al. 2012; Stevison et al. 2016; Pfeifer 2020a; Soni et al. 2025a; Versoza et al. 2025), we performed further filtering to improve the precision of the variant call set in order to reduce potential spurious recombination events. Namely, we excluded SNPs if sites were (1) located within a cluster of variants (defined as more than 3 SNPs within a 10 bp window; calculated using GATK’s *VariantFiltration* with the “*--cluster-size* 3 *--cluster-window-size* 10” parameters), (2) exhibiting an excess of heterozygosity (defined as a Hardy-Weinberg equilibrium *P*-value < 0.01; calculated using VCFtools with the “*--hardy*” option), or (3) located within 5 bp of an insertion/deletion. Additionally, we required that SNPs reciprocally lifted over between the rhesus macaque (rheMac10) and the human (hg38) coordinate systems using the UCSC liftOver tool (Casper et al. 2026), excluding any sites overlapping with the UCSC Problematic Regions track (which contains sub-tracks for UCSC Unusual Regions, ENCODE Blacklist, and GRC Exclusions). To account for regions with poor short-read alignment that can lead to false positive variant calls, we applied a depth of coverage filter, excluding any sites residing in the 2.5% and 97.5% percentiles of the genome-wide distribution (i.e., removing any sites with a read depth *DP* < 242 or *DP* > 400). Afterward, we re-genotyped the filtered dataset using the pan-genome approach implemented in Graphtyper 2.7.2 to improve genotyping accuracy, keeping only sites where alternate alleles exhibited an *AAScore* > 0.5 in the population, as recommended by the developers (Eggertson et al. 2017, 2019). Lastly, we phased variant sites in each population separately using Beagle v.5.5 (Browning et al. 2021). This produced final population-scale datasets of 9.7 million phased SNPs in the ten rhesus macaques of Chinese descent and 13.9 million phased SNPs in the ten rhesus macaques of Indian descent. These two datasets were used to infer population-specific landscapes of recombination (see the section entitled “Inferring fine-scale rates of recombination”).

### Inferring fine-scale rates of neutral divergence and mutation

We indirectly estimated mutation rates from species-level divergence data (see the reviews of Arbogast et al. 2002; Pfeifer 2020b), relying upon the observation that the neutral mutation rate is equivalent to the neutral substitution rate (Kimura 1968). To infer fine-scale neutral divergence, and thus mutation, we initially extracted a subset of the 447-way multi-species alignment, which comprises the mammalian multiple species alignment from the Zoonomia Consortium (Zoonomia Consortium et al. 2020) along with the primate multiple species alignment from the recent study of Kuderna et al. (2024). Specifically, we created two separate subsets of multi-species alignments using the *cactus-hal2maf* function in Cactus v.2.9.8 (Armstrong et al. 2020), including rhesus macaque (*M. mulatta*), either crab-eating macaque (*M. fascicularis*) or human (*H. sapiens*), and a reconstructed ancestor of the two species (i.e., PrimatesAnc133 for *M. mulatta*–*M. fascicularis* and PrimatesAnc007 for *M. mulatta*–*H. sapiens*). For the subset containing *H. sapiens*, we converted the resulting .maf format to .hal format using the *maf2hal* function in Cactus. For the subset including *M. fascicularis*, we first removed the outdated crab-eating macaque assembly using the *halRemoveGenome* function in Cactus and then replaced it with the higher quality telomere-to-telomere assembly now available for the species, T2T-MFA8v1.1 (NCBI RefSeq assembly: GCF_037993035.2; Zhang et al. 2025), by realigning the new *M. fascicularis* assembly to the subset including *M. mulatta* and the reconstructed ancestor (PrimatesAnc133) with Cactus. We then used the resulting .hal files of each subset as input for the *halSnps* function in Cactus, which computes point mutations between a pair of genomes. Specifically, we ran *halSnps* specifying *M. mulatta* as the target species, and either *M. fascicularis* or *H. sapiens* and the respective reconstructed ancestor as our comparison, with the “*--tsv*” flag set to produce output in .bed format. Afterward, we filtered point mutations to only include those where *M. mulatta* contained a different allele than both its neighbor (*M. fascicularis* or *H. sapiens*) and the reconstructed ancestor, ensuring that substitutions occurred on the *mulatta* branch. To guard against polymorphisms, any sites known to segregate in either rhesus macaques (see the section entitled “Short-read data”) or, in the *H. sapiens* comparison, humans (based on data from the Yoruban population included in the 1000 Genomes Project; 1000 Genomes Project Consortium 2015) were removed from the dataset (note that no equivalent dataset was available for crab-eating macaques). To this end, we lifted putative substitutions on the *mulatta* branch over to the human coordinate system (hg38) and then back to rhesus macaque (rheMac10) using the *halLiftover* function in Cactus, keeping only those that lifted back to their original position; similarly, we lifted SNPs known to segregate in humans over to the rhesus macaque coordinate system.

In order to ensure that the final dataset contained neutral substitutions within high-quality regions of the multi-species alignments, we used the Biopython tool suite (Cock et al. 2009) to extract subsets from the .maf files in a way such that individual alignment blocks (1) contained information for *M. mulatta*, either *M. fascicularis* or *H. sapiens*, and the reconstructed ancestor of the two species, without any gaps (-) or missing values (N), (2) were accessible to the long-read data of our study (see the section entitled “Long-read data”), (3) were located outside of repetitive regions (as annotated in the rheMac10 genome assembly; Warren et al. 2020) which are prone to spurious variant calls from short-read data (Pfeifer 2017), and (4) were located more than 10 kb away from exons (as annotated in the rheMac10 genome assembly; Warren et al. 2020) or constrained sequence elements (reported by Kuderna et al. 2024) to avoid the biasing effects of selection at linked sites. We then separated the resulting subsets into sliding windows of 1 kb, 10 kb, 100 kb, and 1 Mb with 500 bp, 5 kb, 50 kb, and 500 kb step sizes, respectively, excluding any windows for which < 75% of sites were accessible. We calculated divergence by taking the number of neutral substitutions per window divided by the number of accessible sites per window. Finally, we converted divergence rates to mutation rates by dividing the divergence estimate by the estimated divergence times in *M. mulatta* generations, assuming divergence times of 25 million years ago (mya) and 35 mya with *H. sapiens* (Goodman et al. 1998; and see the reviews by Disotell and Tosi 2007; Skipper 2007; Chintalapati and Moorjani 2020), 2.7 mya and 4.2 mya with *M. fascicularis* (Bergeron et al. 2021; Tan et al. 2023), and *M. mulatta* generation times of 9 years and 11 years (Gagliardi et al. 2007; Xue et al. 2016).

### Inferring fine-scale rates of recombination

We applied two commonly used methods to infer the recombination landscapes in the two populations of rhesus macaque: LDhat v.2.2 (Auton and McVean 2007) and pyrho v.0.1.7 (Kamm et al. 2016; Spence and Song 2019). Both methods rely on patterns of LD to infer recombination rates, however only pyrho is able to directly account for the demographic history of a population. We used both short- and long-read datasets for the Chinese and Indian rhesus macaque populations as input for this recombination inference (see the sections entitled “Short-read data” and “Long-read data” above for details), following previously described methodology in non-human primates (Auton et al. 2012; Stevison et al. 2016; Pfeifer 2020a).

For LDhat, we first generated likelihood lookup tables for ten diploid individuals (*-n* 20, i.e., 20 haplotypes) in each dataset using the *complete* function in LDhat with the population-specific estimates of genetic diversity, *θ_W_* (*-theta*) (Watterson 1975), a maximum population-scaled recombination rate, *ρ*, of 100 (*-rhomax* 100), and 201 grid points (-*n_pts* 201) for inference. We then pre-processed the polymorphism data by first splitting each chromosome in each dataset into segments of 4,000 SNPs with a 200 SNP overlap between adjacent regions and then generating the required LDhat .sites and .locs input files using the “*--ldhat*” function in VCFtools v.0.1.14 (Danecek et al. 2011). As singletons can reduce the power to accurately infer recombination hotspots in populations which have experienced size expansions (Dapper and Payseur 2018) — as is the case in the Chinese rhesus macaque population (Heenkenda et al. 2026) — recombination rate inference was performed both with and without singletons for comparison. Specifically, based on these likelihood lookup tables and polymorphism data, we inferred population-specific fine-scale recombination rate maps using the *interval* function in LDhat with a block penalty of 5 (*-bpen* 5), and ran 60 million iterations (*-its* 6000000) with a sampling scheme of 40,000 iterations (*-samp* 40000). Afterward, we discarded the burn-in (here the first 20 million iterations; *-burn* 500) of the Monte Carlo Markov Chain using the *stat* function in LDhat. We then merged the resulting region-based recombination rate estimates, keeping overlapping region estimates with the smallest confidence intervals. In order to account for incorrectly called or genotyped variants that are likely to interrupt genuine patterns of LD (Auton et al. 2012; Pfeifer 2020a), we identified regions where population-scaled estimates were ≥ 100 *ρ*/kb or where gaps > 50 kb were observed in the reference assembly, and set the recombination rates of these problematic regions, as well as those of the surrounding 100 SNPs (i.e., 50 SNPs up- and down-stream), to 0 (Table S1). As LDhat produces population-scaled recombination rates, we used the effective population sizes (*N_e_*) to convert *ρ* to *r*. To this end, we calculated *N_e_* for each population as *θ_W_ / 4μ*, assuming a mutation rate, *μ*, of 0.58 × 10^-8^ /bp/gen (i.e., the pedigree-based estimate obtained by Wang et al. [2020]) or 1.49 × 10^-8^ /bp/gen (i.e., the maximum indirect estimate observed from patterns of *M. mulatta*–*H. sapiens* divergence in this study; see the section entitled “Inferring fine-scale rates of neutral divergence and mutation”).

Unlike LDhat, pyrho can take as input an inferred demographic history when estimating recombination rates. As such, we inferred fine-scale recombination maps using pyrho together with the demographic histories recently inferred by Heenkenda et al. (2026) for these two populations. In brief, Heenkenda et al. (2026) considered two mutation rate regimes in their models: a “low-mutation rate” regime based on the pedigree-based estimate of Wang et al. (2020) (*μ* = 0.58 × 10^-8^ /bp/gen) and a “high-mutation rate” regime based on the maximum indirect estimate inferred in this study from the patterns of *M. mulatta*–*H. sapiens* divergence (*μ* = 1.49 × 10^-8^ /bp/gen). Under the “low-mutation rate” regime, the Chinese and Indian rhesus macaque populations split from the ancestral population (*N_anc_* = 64,723) approximately 139,371 generations ago, resulting in population sizes of 76,025 and 270,072 individuals, respectively. Around 71,781 generations ago, the Chinese population began to expand, reaching a final population size of 219,783 individuals. In contrast, the Indian population experienced a population bottleneck around 8,814 generations ago, resulting in a final population of 14,091 individuals. Migration rates under this demographic history were 6.63 × 10^-6^ individuals per generation from the Chinese to the Indian population and 1.46 × 10^-5^ individuals per generation from the Indian to the Chinese population. Under the “high-mutation rate” regime re-scaling, the Chinese and Indian rhesus macaque populations split from the ancestral population (*N_anc_* = 21,986) approximately 80,351 generations ago, resulting in population sizes of 24,710 and 45,917 individuals, respectively. Around 31,538 generations ago, the Chinese population began to expand, reaching a final population size of 97,535 individuals. The Indian population experienced a population bottleneck around 2,884 generations ago, resulting in a final population of 5,601 individuals. Migration rates under this demographic history were 1.16 × 10^-5^ individuals per generation from the Chinese to the Indian population and 4.10 × 10^-5^ individuals per generation from the Indian to the Chinese population.

For pyrho, we first generated likelihood lookup tables using the *make_table* function, specifying population sizes (*--popsizes*) and timing of population size changes (*--epochtimes*) corresponding to either the Chinese or Indian population under each of the two demographic scenarios described above. Next, we used the *hyperparam* function to determine the optimal block penalty and window size for each run, specifying the mutation rate corresponding to the demographic model being tested. Based on the optimal block penalty and window size observed for each population-level dataset and each demographic model, we inferred population-specific fine-scale recombination rate maps using the *optimize* function. As with LDhat, we set the recombination rates of regions where population-scaled estimates were ≥ 100 *ρ*/kb or where gaps > 50 kb were observed in the reference assembly to 0 (Table S1).

In order to convert recombination rates inferred by LDhat and pyrho to units of physical genomic distance, we re-scaled rates to the autosomal genetic map length observed from pedigree data (2357.1 cM; Versoza et al. 2024). We calculated mean recombination rates in sliding windows of 1 kb, 10 kb, 100 kb, and 1 Mb with 500 bp, 5 kb, 50 kb, and 500 kb step sizes, respectively, and computed Pearson correlation coefficients between the maps using the *cor()* function implemented in the *pearson* package in R v.4.5.1 (R Core Team 2025).

### Assessing the performance of LDhat and pyrho under the demographic histories of the Chinese and Indian rhesus macaque populations

To assess the performance of LDhat and pyrho under the demographic histories inferred for the Chinese and Indian rhesus macaque populations, we simulated 1 Mb genomic regions for ten individuals from each population in msprime v.1.3.4 (Baumdicker et al. 2022). Specifically, we simulated 10 replicates under the demographic models of Heenkenda et al. (2026) — consisting of a split of the ancestral population into Chinese and Indian populations, two size changes per population, and asymmetric migration between the two populations (as described in the section “Inferring fine-scale rates of recombination”) — assuming a mutation rate of either 0.58 × 10^-8^ /bp/gen (based on the pedigree-based estimate of Wang et al. [2020]) or 1.49 × 10^-8^ /bp/gen (based on the maximum indirect estimate inferred in this study from the patterns of *M. mulatta*–*H. sapiens* divergence) and a recombination rate of 0.88 cM/Mb (as inferred by Versoza et al. [2024] from pedigree data). We estimated recombination rates from this simulated data using LDhat and pyrho in order to assess any underlying bias arising from these rhesus macaque-specific demographic histories.

### Inferring recombination hotspots

We identified recombination hotspots harbored in the Chinese and Indian rhesus macaque populations following the methodology described in Hoge et al. (2024). We first split each chromosome into 2kb-windows overlapping by 1 kb and then calculated the average recombination rate per window for each population based on the long-read maps inferred by LDhat. For each window containing > 50% accessible sites (defined here as sites accessible to our long-read data where population-scaled estimates were < 100 *ρ*/kb and where no gaps > 50 kb were observed in the reference assembly; see the section entitled “Inferring fine-scale rates of recombination”), we calculated its intensity or “heat” by dividing the mean recombination rate in the central window by the mean rate observed in the 20 kb flanking regions surrounding the central window (i.e., the genomic background). We defined candidate hotspots as windows with a heat exceeding 5· or 10· the genomic background, and subsequently merged candidate windows located within 5 kb of each other to produce the final sets of population-specific hotspots. Based on these datasets, we identified overlaps between the 10× hotspots in the Chinese and Indian populations using the *GenomicRanges* package (Lawrence et al. 2013) in R v.4.5.1 (R Core Team 2025).

Because the locations of recombination hotspots are thought to be primarily controlled by the sequence-specific binding of PRDM9 in primates (see the review by Paigen and Petkov 2018), we leveraged the published protein sequence of PRDM9 in rhesus macaques (NCBI Reference Sequence: XP_028705194.2; Warren et al. 2020) to predict a position weight matrix for the DNA binding specificity of the C_2_H_2_ zinc fingers using the https://zf.princeton.edu webserver under the polynomial support vector machine model described in Persikov et al. (2009). Given that three to four sequential zinc finger domains are generally required to specifically bind DNA and given that primate PRDM9 binding motifs are often 13–15 bp long (e.g., Myers et al. 2008; Auton et al. 2012; Schwartz et al. 2014), we used the FIMO function (Grant et al. 2011) of the MEME software suite v.5.5.9 (Bailey et al. 2015) to scan the 5× hotspots for overlaps of putative binding motifs based on sliding windows of 13, 14, and 15 bp along the position weight matrix and a background model generated from the rhesus macaque genome assembly using the *nuc* function in BEDTools v.2.31.1 (Quinlan 2014), applying a *P*-value cut-off of 0.05. As the published protein sequence of PRDM9 was obtained from a single individual, we additionally leveraged high-resolution crossover events (i.e., crossovers with a resolution < 10 kb) previously observed in pedigrees (Versoza et al. 2024) to expand our search for putative PRDM9 binding motifs. To this end, we used the ZOOPs model in MEME v.5.5.8 (Bailey and Elkan 1994) together with 65 high-resolution crossover breakpoints — each extended by 500 bp to increase the likelihood of capturing the complete region subject to the double-strand break and repair machinery — and a genomic background that included randomly selected non-breakpoint regions with similar genomic characteristics (i.e., regions of the same length with a GC-content within 5% of the matched breakpoint region) to account for possible nucleotide sequence bias, setting the discovery threshold of the analysis to 1E-5. To compare the distribution of putative binding motifs in breakpoint regions to those in non-breakpoint regions, we performed permutation tests using MOODS v.1.9.4.1 (Korhonen et al. 2009) along with an additional 1,000 random non-breakpoint regions, requiring a *P*-value < 0.05 for a match. To test for differences between breakpoint and non-breakpoint regions, we performed a Fisher’s exact test in R v.4.2.2 (R Core Team 2025).

In order to determine whether the enrichment of a putative binding motif within 5× hotspots was statistically significant, we created a dataset of sequence-matched cold spots. Specifically, following Hoge et al. (2024), we matched each hotspot with a randomly selected cold spot of the same length that exhibited a similar GC-content (no more than 2.5% deviation) and a mean recombination rate less than half of the genome-wide average. Additionally, we required that each cold spot was located no farther than 5 Mb from its matched hotspot and that it was not in the immediate vicinity (within 10 kb) of another hotspot or cold spot. To assess enrichment, we used a binomial test to compare the proportion of hotspots containing a putative binding motif with the proportion expected from the sequence-matched cold spots. Finally, we used AlphaFold 3 (Abramson et al. 2024) to characterize the binding of the best putative binding motif and the rhesus macaque PRDM9 sequence *in silico*.

### Correlations of rates of recombination with genomic features

To better understand how rates of recombination correlate with genomic features, we calculated a variety of summary statistics for each population across 10 kb, 100 kb, and 1 Mb sliding windows with 5 kb, 50 kb, and 500 kb step sizes, respectively. These summary statistics included: recombination rate (based on the long-read maps inferred by LDhat; see the section entitled “Inferring fine-scale rates of recombination”), divergence from both *H. sapiens* and *M. fascicularis* (see the section entitled “Inferring fine-scale rates of neutral divergence and mutation”), repeat content and gene content (both based on the annotations available for the rheMac10 genome assembly; Warren et al. 2020). In addition, we calculated per-site nucleotide diversity based on the long-read data (see the section entitled “Long-read data”) using the “*--stats pi*” function in pixy v.2.0.0.beta14 (Korunes and Samuk 2021). We identified CpG-islands using the *cpgplot* function implemented in EMBOSS v.6.6.0 (Rice et al. 2000) together with the rhesus macaque genome assembly and analyzed GC-content using the *nuc* function in BEDTools v.2.31.1 (Quinlan 2014). We performed Spearman rank correlations using the *cor_mat()* function from the rstatix package (Kassambara 2023) in R v.4.5.1 (R Core Team 2025).

## Results and Discussion

### Fine-scale rates of neutral divergence and mutation

In order to estimate fine-scale mutation rates across the rhesus macaque genome, we calculated neutral divergence from comparisons with high-quality genomes of a close relative, the crab-eating macaque (*Macaca fascicularis*), as well as from a more distant relative, humans (*Homo sapiens*). We masked both non-neutral regions as well as any sites segregating within the species from the comparative datasets before identifying substitutions between *M. mulatta*, either *M*. *fascicularis* or *H. sapiens,* and their respective reconstructed ancestor. We leveraged neutral substitutions on the *M. mulatta* branch to calculate neutral divergence relative to the number of accessible sites in sliding windows of 1 kb, 10 kb, 100 kb, and 1 Mb regions across the rhesus macaque genome (see “Materials and Methods” for details). The average genome-wide neutral divergence from *M. fascicularis* varied from 0.001468 (±) 0.001855 in 1 kb windows to 0.001521 (±) 0.000259 in 1 Mb windows, while from *H. sapiens*, divergence varied from 0.032420 (±) 0.010407 in 1 kb windows to 0.033898 (±) 0.003486 in 1 Mb windows (Table S2).

We converted these divergence rates to mutation rates by dividing the divergence estimates by the estimated divergence times in *M. mulatta* generations. As for many species, generation time estimates in rhesus macaque populations contain some uncertainty given that this parameter is not directly observable and variation exists with regards to reproductive life cycles; similarly, uncertainty exists in estimates of divergence times between *M. mulatta* with both *M. fascicularis* and *H. sapiens* due to incomplete fossil records and limitations inherent to the calibration of the “molecular clock”. Therefore, to calculate fine-scale mutation rates, we considered well-supported generation times of both 9 years and 11 years (Gagliardi et al. 2007; Xue et al. 2016) as well as divergence times of 2.7 mya and 4.2 mya for *M. mulatta–M. fascicularis* (Bergeron et al. 2021; Tan et al. 2023) and divergence times ranging from 25 mya to 35 mya for *M. mulatta–H. sapiens* (Goodman et al. 1998; and see the reviews by Disotell and Tosi 2007; Skipper 2007; Chintalapati and Moorjani 2020). Calculating genome-wide neutral mutation rates in sliding genomic windows from divergence with *M. fascicularis*, estimates ranged between 0.32 × 10^-8^ /bp/gen based on a generation time of 9 years and a divergence time of 4.2 mya, and 0.62 × 10^-8^ /bp/gen based on a generation time of 11 years and a divergence time of 2.7 mya (see Figure 1A for the genome-wide distribution of mutation rates at 10 kb, Figure S2 for the corresponding fine-scale mutation rate maps, and Table S3 for a summary of the mutation rate estimates at different window sizes). Notably, these estimates are in line with direct observations from rhesus macaque pedigrees, with Wang and colleagues (2020) estimating a mean per-site per-generation mutation rate of 0.58 × 10^-8^ based on 14 parent-offspring trios with an average parental age of 7.5 years (range: 0.23 × 10^-8^ to 1.18 × 10^-8^). More recent estimates by Bergeron et al. (2021) based on 19 parent-offspring trios were slightly higher, reporting a mean per-site per-generation mutation rate of 0.77 × 10^-8^ (range: 0.49 × 10^-8^ to 1.16 × 10^-8^), though the average parental age of the individuals included in their study was older as well (10.4 years). Mutation rate estimates based on divergence with *H. sapiens* were substantially higher than those based on divergence with *M. fascicularis*, ranging between 0.83 × 10^-8^ /bp/gen based on a generation time of 9 years and a divergence time of 35 mya, and 1.49 × 10^-8^ /bp/gen based on a generation time of 11 years and a divergence time of 25 mya (see Figure 1B for the genome-wide distribution of mutation rates at 10 kb, Figure S3 for the corresponding fine-scale mutation rate maps, and Table S4 for a summary of the mutation rate estimates at different window sizes). These estimates are more similar to *de novo* point mutation rates previously observed in humans and other great apes, which range between 0.97 × 10^-8^ /bp/gen and 1.66 × 10^-8^ /bp/gen (Conrad et al. 2011; Venn et al. 2012; Besenbacher et al. 2019; and see Bergeron et al. 2023 and references therein). Taken together, our results are thus consistent with higher per-generation mutation rates along the hominoid branch (Bergeron et al. 2023), while the *M. mulatta* branch is likely better characterized by the reduced rates inferred in comparison with *M. fascicularis*.

**Figure 1.**
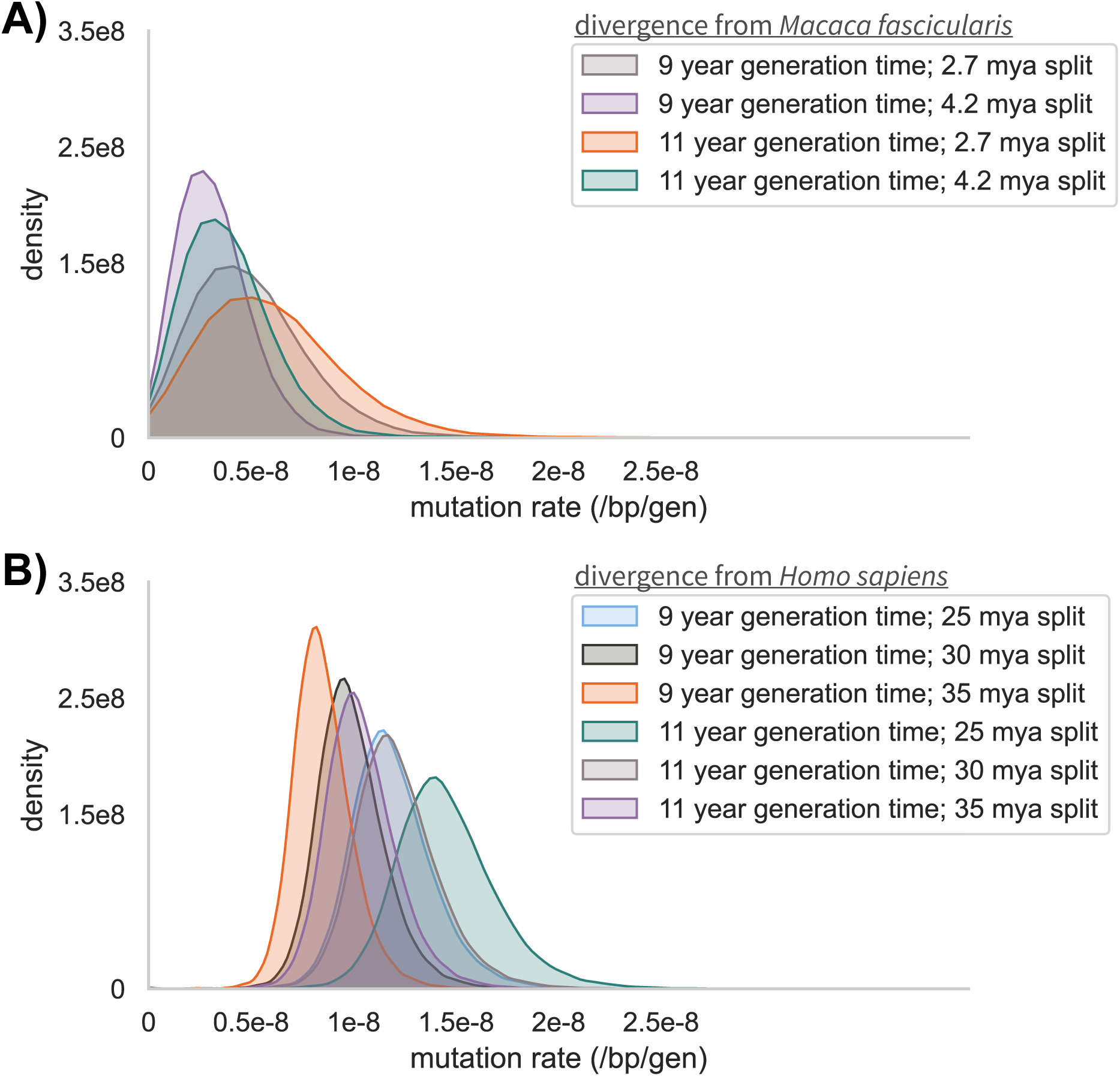
Frequency distribution of fine-scale per-site per-generation mutation rates based on the mean neutral divergence from (a) *M. fascicularis* and (b) *H. sapiens* calculated in sliding windows of 10 kb across the genome, assuming generation times of 9 years and 11 years (Gagliardi et al. 2007; Xue et al. 2016) as well as divergence times for *M. mulatta–M. fascicularis* of 2.7 mya and 4.2 mya (Bergeron et al. 2021; Tan et al. 2023) and divergence times for *M. mulatta–H. sapiens* ranging from 25 mya to 35 mya (Goodman et al. 1998; Disotell and Tosi 2007; Skipper 2007; Chintalapati and Moorjani 2020).

### Fine-scale rates of recombination

As one of the most important primate models in biomedical research, genetic linkage maps have been generated to facilitate genomics research using both human microsatellite loci (Rogers et al. 2006) and pedigree data (Versoza et al. 2024). Although these maps offer key insights into the rates of recombination at the coarse-scale, they cannot capture the heterogeneity in recombination at the fine-scale — knowledge that is essential for many genetic analyses, including genome-wide association studies and scans for the targets of selection. Thus, in order to obtain such population-specific fine-scale maps for rhesus macaques of Chinese and Indian descent, we applied two commonly used methods, LDhat (Auton and McVean 2007) and pyrho (Kamm et al. 2016; Spence and Song 2019), both of which infer fine-scale recombination rates from patterns of LD. Despite measuring the same underlying process, LD-based recombination rate estimates often differ from pedigree-based rates for a variety of reasons. For example, while pedigree-based linkage maps directly measure the rate of crossover at the time of sampling, indirect LD-based maps reflect historically-averaged recombination rates which can be sensitive to other evolutionary processes such as the population-specific demographic history (including changes in population size and rates of gene flow), selective pressures over time, and changes in hotspot activity (e.g., due to PRDM9 binding site turnover; Schwartz et al. 2014; Stevison et al. 2016; Dapper and Payseur 2018; Samuk and Noor 2022; Dutheil 2024).

Given the sensitivity of LD-based approaches to long-term evolutionary processes, we first quantified the performance of both methods using simulated datasets of the Chinese and Indian population histories. For this testing, replicates of 1 Mb genomic regions were simulated following the previously inferred demographic histories of the two populations, including split times, population size changes, and migration rates (Heenkenda et al. 2026), assuming a constant recombination rate (0.88 cM/Mb; as inferred by Versoza et al. [2024] from pedigrees) as well as a constant mutation rate of either 0.58 × 10^-8^ /bp/gen (i.e., the mean pedigree-based estimate of Wang et al. [2020]) or 1.49 × 10^-8^ /bp/gen (i.e., the maximum indirect estimate observed from patterns of *M. mulatta*–*H. sapiens* divergence in this study), drawing 10 Chinese and 10 Indian individuals per replicate to match our empirical sample size (see “Materials and Methods” for details). Results suggest that LDhat — which is naïve of the underlying population histories — generally underestimates recombination rates, with mis-inference more pronounced in the Indian population (Figure S4) which has experienced a severe population bottleneck ∼3,000–9,000 generations ago (Heenkenda et al. 2026). In contrast, despite incorporating the inferred demographic histories as input, pyrho tends to overestimate recombination rates under these population-specific histories, again, particularly for the Indian population (Figure S4). These observations are in broad agreement with previous simulation-based performance benchmarks in showing that recombination rates are often underestimated in bottlenecked populations when demography is neglected (as is the case for LDhat) but can also be overestimated even when demographic effects are explicitly modeled (as is the case for pyrho), particularly when sample sizes are small (≤ 50 individuals; Dutheil 2024). In primates specifically, Soni et al. (2025a) found that pyrho underestimated recombination rates in aye-ayes, a critically endangered strepsirrhine characterized by historical bottlenecks and a small *N_e_* (Terbot et al. 2025b); similarly, a pattern of underestimation was recently reported for both LDhat and pyrho in common marmosets (Soni et al. 2025b), a platyrrhine characterized by a strong population decline ∼7,000 years ago (Soni et al. 2025c). Additionally, the neglect of gene conversion — an alternative outcome of recombination that occurs at higher rates than crossovers in primates and that can have a particularly strong impact in populations of declining size (Williams et al. 2015; Halldorsson et al. 2016; Wall et al. 2022; Dutheil 2024; Versoza et al. 2024, 2025; Palsson et al. 2025) — likely contributes to the observed mis-inference. Moreover, previous research has demonstrated that recombination rates may show under- or over-estimation in populations characterized by significant migration (*N_e_m* ≥ 1), depending on the timing and duration of gene flow (Samuk and Noor 2022). Given that conservative estimates of *Nₑm* for the Chinese and Indian populations range from 4.2 to 7.3 individuals per generation based on the smallest migration rate inferred from the demographic models (Heenkenda et al. 2026), gene flow thus likely further contributes to the uncertainty in recombination rate estimates.

The Chinese rhesus macaque population was characterized by higher empirical estimates of recombination rate inferred using LDhat compared to those inferred with pyrho; however, this pattern was reversed for the Indian population (see Figure 2 for per-site per-generation recombination rate estimates at the 1Mb-scale and Figures S5 and S6 for 10kb- and 100kb-scales, respectively), suggesting that method choice introduces systematic differences depending on the population-specific history. Despite these differences, rate estimates were overall relatively consistent between the two methods for the Chinese population, particularly when based on the long-read data; in contrast, rates inferred for the Indian population exhibited much larger discrepancies (see Table S5 for average per-site per-generation recombination rate estimates based on 1 kb, 10 kb, 100 kb, and 1 Mb windows across the genome). Given that the demographic history of the Indian rhesus population includes a more severe population bottleneck, an underestimation of recombination rates by LDhat is consistent with expectations based on both previous work (Dapper and Payseur 2018; Dutheil 2024) as well as our simulation results (Figure S4); similarly, in agreement with our simulations indicating that pyrho tends to overestimate recombination rates under these population histories, the higher estimates observed here are likely to be expected. In both populations, the estimates based on the higher-quality long-read data were consistently lower than those based on the short-read data (Figure 2; and see Table S5) as errors common in the statistical phasing of short-read data can increase recombination rate estimates by introducing spurious breaks in the patterns of LD. Specifically, based on the long-read data and assuming a mutation rate of 1.49 × 10^-8^ /bp/gen, average per-site per-generation recombination rate estimates ranged from 1.45 × 10^-8^ (pyrho) to 1.70 × 10^-8^ (LDhat) in the Chinese population and from 0.40 × 10^-8^ (LDhat) to 1.92 × 10^-8^ (pyrho) in the Indian population. In contrast, using short-read data, rate estimates ranged from 1.71 × 10^-8^ (pyrho) to 2.58 × 10^-8^ (LDhat) in the Chinese population and from 0.53 × 10^-8^ (LDhat) to 3.17 × 10^-8^ (pyrho) in the Indian population. Under the demographic model with *μ* = 0.58 × 10^-8^ /bp/gen, recombination rates were considerably lower, ranging from 0.60 × 10^-8^ (pyrho) to 0.66 × 10^-8^ (LDhat) based on long-read data in the Chinese population (short-read data: 0.71 × 10^-8^ [pyrho] to 1.00 × 10^-8^ [LDhat]) and from 0.16 × 10^-8^ (LDhat) to 0.84 × 10^-8^ (pyrho) based on long-read data in the Indian population (short-read data: 0.21 × 10^-8^ [LDhat] to 1.40 × 10^-8^ [pyrho]), owing to the larger *N_e_* of the model (and thus an accompanying reduction in *r*). Including singletons, recombination rates were slightly lower across both mutation rates and populations, particularly within the Chinese population. Specifically, based on the long-read data, average per-site per-generation recombination rate estimates within the Chinese population varied across both applied mutation rates from 0.54 × 10^-8^ to 1.52 × 10^-8^ /bp/gen and from 0.15 × 10^-8^ to 0.42 × 10^-8^ /bp/gen within the Indian population using LDhat (Table S6). Using the same dataset but excluding singletons, recombination rates estimated with LDhat within the Chinese population varied across both applied mutation rates from 0.63 × 10^-8^ to 1.70 × 10^-8^ /bp/gen and from 0.16 × 10^-8^ to 0.42 × 10^-8^ /bp/gen within the Indian population (Table S5). This is consistent with previous results suggesting that the inclusion of singletons can reduce power to accurately infer recombination, particularly in populations having experienced a recent expansion (Dapper and Payseur 2018).

**Figure 2.**
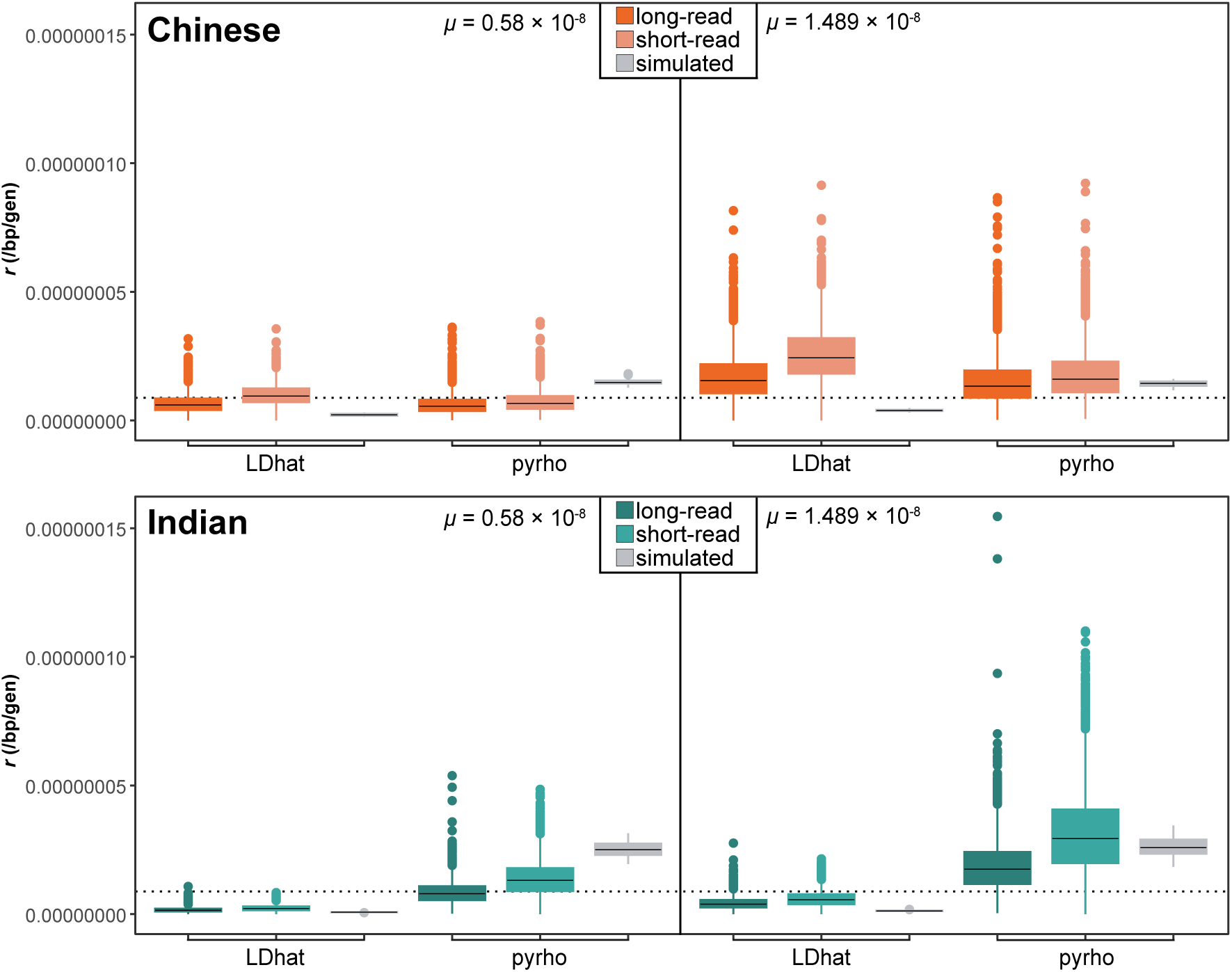
Per-site per-generation recombination rate estimates for rhesus macaque populations of (top) Chinese descent and (bottom) Indian descent as inferred using LDhat and pyrho at the 1Mb-scale across the genome. Estimates obtained from empirical long-read and short-read data are shown in dark and light colors, respectively (orange: Chinese; teal: Indian). For comparison, estimates obtained from simulations under the population-specific demographic histories recently inferred by Heenkenda et al. (2026) assuming a constant recombination rate (0.88 cM/Mb; as observed in pedigrees by Versoza et al. [2024]; indicated by a dotted line) are shown in gray. Population-scaled recombination rate estimates were converted to per-generation estimates using population-specific *N_e_* values based on mutation rates, *μ*, of 0.58 · 10^-8^ /bp/gen (i.e., the pedigree-based estimate obtained by Wang et al. [2020]) or 1.49 · 10^-8^ /bp/gen (i.e., the maximum indirect estimate observed from patterns of *M. mulatta*–*H. sapiens* divergence in this study).

Because LD-based recombination rate estimators tend to systematically over- or under-estimate rates as shown above, it is often useful to scale indirectly inferred rates to a pedigree-derived genetic linkage map, thereby correcting the overall scale while preserving rate heterogeneity across the map. Following previous work in non-human primates (Wall et al. 2022; Soni et al. 2025a, 2026), per-site per-generation recombination rates were thus re-scaled to the genetic map length previously obtained from pedigree data (2357.1 cM; Versoza et al. 2024). Across both methods and datasets, re-scaled average recombination rates varied from 0.909 (±) 0.111 to 0.925 (±) 0.163 cM/Mb in the Chinese population, and from 0.904 (±) 0.131 to 0.915 (±) 0.158 cM/Mb in the Indian population (Table S7). As observed in other species (Zedek et al. 2026), chromosome length was positively associated with recombination rate, with estimated rates ranging from ∼0.8 cM/Mb on the longest autosome (chromosome 1) to ∼1.2 cM/Mb on the shortest autosome (chromosome 19), with heterogeneity observed between the two populations (Figure S7). Comparatively, these re-scaled genome-wide recombination rate estimates from the two rhesus macaque populations are markedly higher than those previously inferred for the species by Xue et al. (2016, 2020) from short-read data and without any consideration of the underlying population history (0.433 ± 0.333 cM/Mb to 0.448 ± 0.286 cM/Mb); indeed, these rates are only slightly lower than those previously inferred in humans (∼1.0 cM/Mb; Palsson et al. 2025) and other great apes (bonobos, chimpanzees, and gorillas: ∼1.2 cM/Mb; Stevison et al. 2016). Moreover, consistent with other primates (Auton et al. 2012; Stevison et al. 2016; Pfeifer 2020a; Wall et al. 2022; Versoza et al. 2024; Soni et al. 2025a,b, 2026), recombination rates in both populations tended to be higher near telomeric regions and reduced around centromeres and adjacent pericentromeric regions (see Figures 3 and S8 for fine-scale recombination rate maps inferred from long-read and short-read data, respectively).

**Figure 3.**
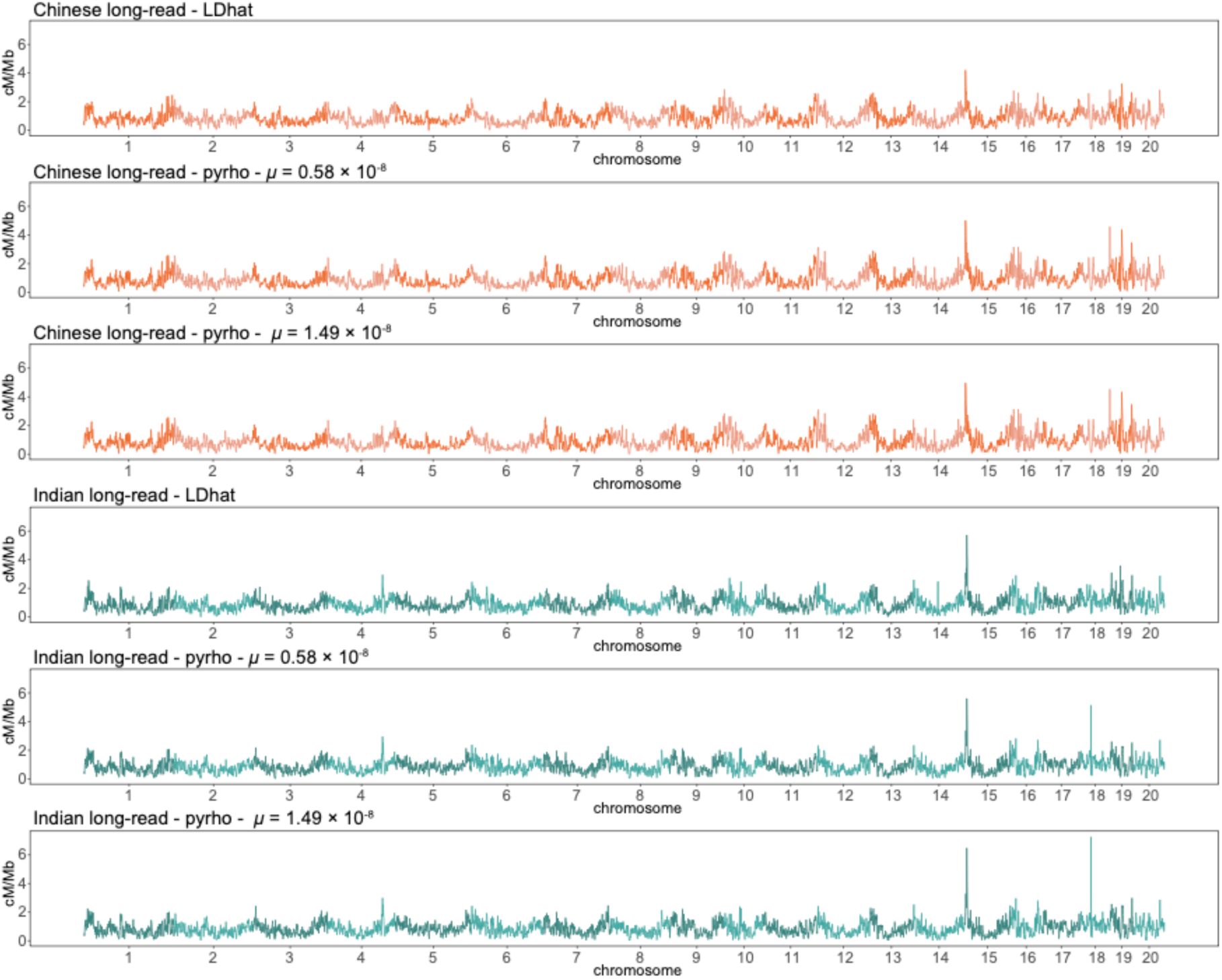
Fine-scale recombination rate maps for rhesus macaque populations of Chinese descent (shown in orange) and Indian descent (teal) as inferred from long-read data using LDhat and pyrho at the 1Mb-scale across the genome. Population-scaled recombination rate estimates were converted to per-generation estimates using population-specific *N_e_* values based on mutation rates (*μ*) of 0.58 · 10^-8^ /bp/gen (i.e., the pedigree-based estimate obtained by Wang et al. [2020]) or 1.49 · 10^-8^ /bp/gen (i.e., the maximum indirect estimate observed from patterns of *M. mulatta*–*H. sapiens* divergence in this study) and re-scaled to the genetic map length previously observed from pedigree data (Versoza et al. 2024). Individual chromosomes are shown in alternating shading.

At the 1Mb-scale, recombination rate maps of Chinese and Indian populations inferred using LDhat and pyrho were strongly positively correlated in both long-read (Pearson’s *r* > 0.72; Figure S9) and short-read datasets (Pearson’s *r* > 0.70; Figure S10), suggesting that broad-scale features of the recombination landscape are conserved between the two populations due to their shared chromosome structure — a pattern similar to that observed in human populations (Serre et al. 2005; Hinch et al. 2011). However, as expected from the rapid evolution of PRDM9 which directs recombination in primates (Myers et al. 2010), this correlation breaks down at the finer scales (with a Pearson’s *r* of *∼*0.2 observed at the 1kb-scale; Figures S9 and S10). Population-specific recombination rate maps inferred in pyrho from the same read type (i.e., long- or short-reads) but under different demographic models were nearly perfectly correlated in the Chinese population (Pearson’s *r* > 0.99 for all scales) (Figure S11); in contrast, in the Indian population, correlations between demographic models were near perfect at the broad (1Mb) scale (Pearson’s *r* > 0.98) but considerably weaker at the finer scales (Pearson’s *r* > 0.89 at the 1kb-scale). The latter owes to the loss in power to detect recombination events as variation decreases, driven here both by the population bottleneck together with the reduced mutation rate model. Recombination rate maps inferred from different read types of the same population were strongly correlated at the broad-scale (with a Pearson’s *r* between 0.73 and 0.87 at the 1Mb-scale; see Figures S12 and S13 for the Chinese and Indian population, respectively) but weakly correlated at the fine-scale (with a Pearson’s *r* between 0.37 and 0.59 at the 1kb-scale).

In addition to genuine biological variation, technical artefacts including potential ascertainment biases in short-read data likely contribute to these observed deviations. In this regard, it is noteworthy that recombination, particularly gene conversion events, lead to DNA mismatches that are preferentially repaired with G or C alleles rather than A or T alleles, resulting in higher recombination rates in GC-rich regions (Williams et al. 2015; Porsborg et al. 2025), which are better resolved with long-read sequencing (Browne et al. 2020). Long-read data also provides more accurate haplotype information compared to the statistical phasing of short-read data for which switch errors are common (Amarasinghe et al. 2020; Browne et al. 2020; De Coster et al. 2021; Espinosa et al. 2024), thus likely further decreasing correlations observed between short-and long-read datasets. Moreover, while the assumption of a constant *N_e_* in LDhat generally seems to produce more stable correlations in the Chinese population, pyrho’s sensitivity to demographic model specification can introduce model-dependent variation, with the strong bottleneck experienced by the Indian population resulting in a particularly high variability. Taken together, our results thus suggest that while broad-scale recombination rates are often highly concordant across methods, datasets, and populations, technical noise driven by sequencing technology can affect recombination rate estimates, particularly at the fine-scales necessary for hotspot detection. As the fine-scale recombination landscapes are expected to vary between populations, the higher resolution afforded by long-read sequencing allows for improved insights into finer scale heterogeneity, enabling detection of population differences.

### Recombination hotspots and PRDM9 motif overlap

Like many mammals, the primate genomes studied to date possess recombination hotspots, which are short regions of the genome (typically ∼1–2 kb in length) with elevated recombination rates (Baudat et al. 2010; Myers et al. 2010; Parvanov et al. 2010). In agreement, ∼80% of recombination in Chinese and Indian rhesus macaques occurred in less than 20% of their genomes based on the LDhat estimates obtained from the long-read data (Figure 4A) — a similar proportion as in humans of African descent and other great apes (∼15-20%; 1000 Genomes Project Consortium 2010; Auton et al. 2012; Stevison et al. 2016). Moreover, similar to humans of European descent (1000 Genomes Project Consortium 2010), the more severely bottlenecked rhesus macaque population of Indian descent exhibited a stronger concentration of recombination in hotspot regions (with ∼14% and ∼19% of recombination occurring in ∼80% of the genome in the populations of Indian and Chinese descent, respectively). Technical challenges, including low haplotype resolution, statistical phasing errors, and SNP ascertainment biases, are thought to complicate the detection of recombination hotspots from short-read data as spurious observations can disrupt haplotypes, thus artificially inflating the LD of the genomic background. This issue becomes more prominent in both expanding populations as higher levels of genetic diversity provide more opportunities for mis-inference as well as in bottlenecked populations as patterns of LD extend over longer distances, further decreasing the contrast between recombination hotspots and the genomic background. In agreement, based on the short-read data, the recombination landscapes of both populations were less concentrated, with ∼80% of recombination occurring in ∼42% of the genome in the more genetically-diverse Chinese population and ∼16% in the less genetically-diverse Indian population.

**Figure 4.**
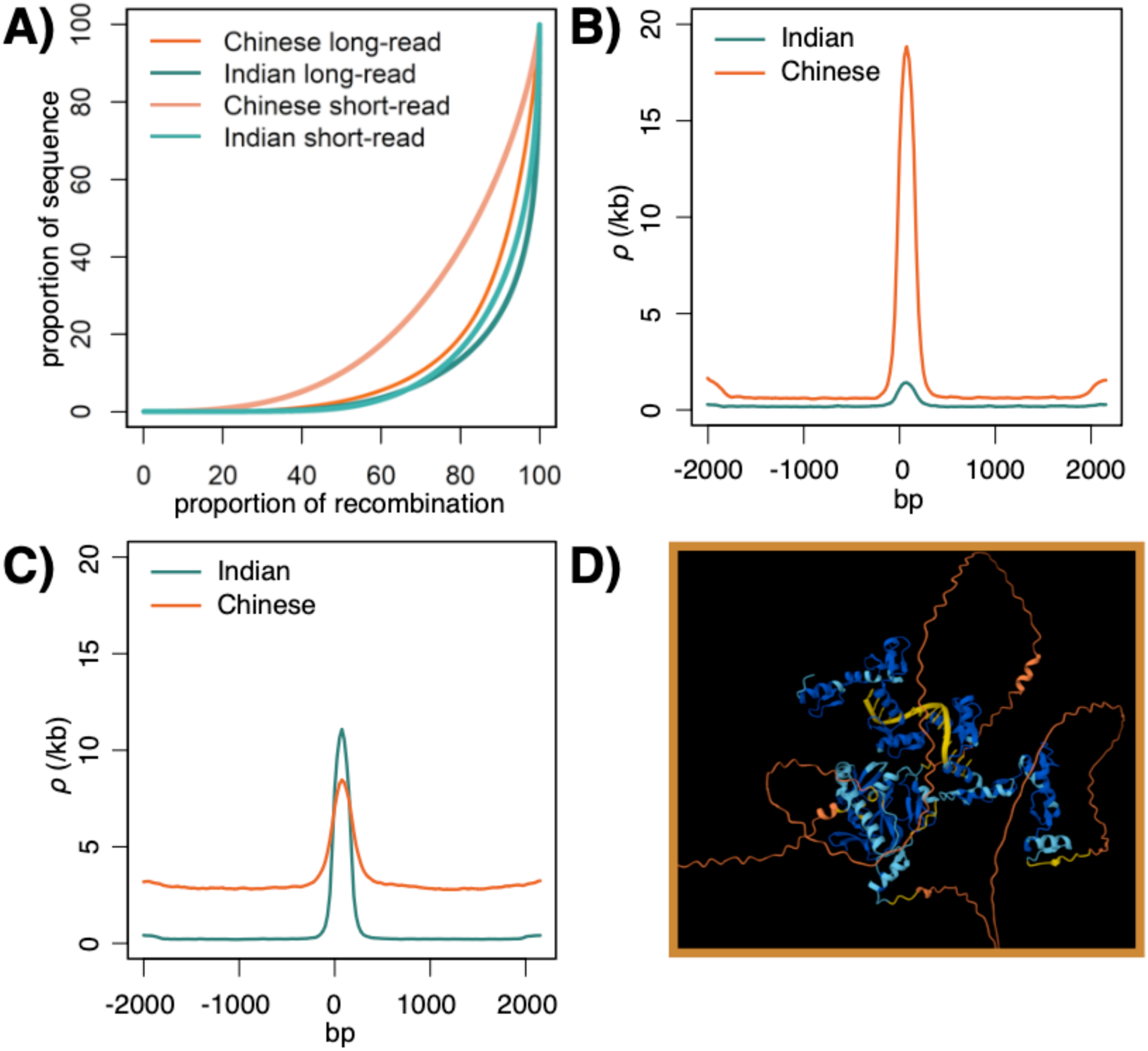
**(A)** Cumulative genome-wide distribution of recombination plotted as the proportion of recombination versus the proportion of sequence for recombination rate maps for rhesus macaque populations of Chinese descent (shown in orange) and Indian descent (teal) as inferred from short-read and long-read data using LDhat. **(B)** Average recombination rates across all hotspots in the Chinese population with a heat exceeding 10× the genomic background. **(C)** Average recombination rates across all hotspots in the Indian population with a heat exceeding 10× the genomic background. **(D)** *In silico* binding prediction of the PRDM9 protein-DNA complex using a sequence from the best predicted PRDM9 binding motif (GGGCTGAGGAGGGGG). Predictions were performed using AlphaFold3 (predictions are provided for non-commercial use only, under and subject to AlphaFold Server Output Terms of Use found under alphafoldserver.com/output-terms).

Leveraging recombination rate estimates based on the higher-quality long-read data, we employed a methodology previously described by Hoge and colleagues (2024) to map recombination hotspots across the genome of both rhesus macaque populations. After filtering to reduce false-positives (see “Materials and Methods” for details), 34,101 (5,700) and 47,024 (25,820) hotspots with a heat exceeding 5· (10·) the genomic background were discovered in the populations of Chinese and Indian descent, respectively (Table S8). Although the number of hotspots is considerably smaller in the Chinese population than in the Indian population, this observation is likely, at least in part, driven by a detection bias due to the differences in population history (see the discussions in Johnston and Cutler 2012; Dapper and Payseur 2018). Specifically, the overall reduction in genome-wide LD combined with an excess of low-frequency variants likely reduces the power to identify localized recombination rate increases in the expanding Chinese population compared to the bottlenecked Indian population. In agreement, using the same dataset but including singletons, even fewer hotspots were detected in the Chinese population: 31,856 and 1,496 hotspots with heat exceeding 5· and 10·, respectively (Table S6). Notwithstanding this, the number of recombination hotspots identified in the two populations is comparable to those previously identified in other primates; for example, while earlier studies in humans have detected roughly 25,000–30,000 recombination hotspots based on both indirect LD-based approaches using short-read data (e.g., International HapMap Consortium 2005,2007; Myers et al. 2005; 1000 Genomes Project Consortium 2010) and direct mapping of double-strand break initiation sites (Pratto et al. 2014), subsequent work providing more complete insights into the landscape of recombination has suggested that the number of hotspots is likely even larger (e.g., Halldorsson et al. 2019). Additionally, the mean width of the hotspots observed in the two rhesus macaque populations (2.3 kb and 2.4 kb for hotspots with a heat exceeding 10× the genomic background in the Chinese and Indian populations, respectively; Table S8) is also reassuringly similar to that observed in human populations (2.3 kb; 1000 Genomes Project Consortium 2010). Notably, although the broad (1Mb) scale recombination landscapes are highly similar, at the fine (hotspot) scale, landscapes exhibit substantial differences between the two populations, with 9,030 (721) of the hotspots exceeding 5· (10·) the genomic background being shared across populations (Figures 4B and 4C; and see Figure S14 for examples of population-specific vs population-shared recombination hotspots). Rhesus macaques thus exhibit a considerably larger proportion of population-specific hotspots (73.5% and 80.8% in the Chinese and Indian populations, respectively), reflecting a high turn-over of hotspot usage driven by variation in PRDM9 alleles, compared to human populations of African and European ancestry (∼10%; Hinch et al. 2011) as might be expected from their higher levels of genetic diversity (∼0.0025 in rhesus macaques vs ∼0.0010 and ∼0.0008 in humans of African and European ancestry, respectively; Xue et al. 2016).

Given that recombination hotspots are primarily controlled by the sequence specific binding of PRDM9 in primates, we sought to predict putative PRDM9 binding motifs and investigate their overlap with recombination hotspots in our datasets. Of the putative binding motifs generated based on the position weight matrix predicted from the published PRDM9 protein sequence, five showed a significant enrichment within recombination hotspots compared to sequence-matched cold spots in the Indian population, with between 36% and 59% of hotspots containing the predicted binding motifs (Table 1). In addition, three putative binding motifs predicted from high-resolution crossover breakpoints observed in pedigrees were significantly enriched within the hotspots of the Indian population, with between 61% and 70% of hotspots containing the predicted binding motifs. In contrast, no significant enrichment of any predicted motifs was observed within the hotspots of the Chinese population — an observation that likely reflects a bias in our study resulting from the use of an Indian-derived reference genome (Warren et al. 2020) as well as crossover breakpoints obtained from a pedigree of Indian rhesus macaques (Versoza et al. 2024) for motif prediction. Together, these factors suggest that PRDM9 binding motifs likely differ between Chinese and Indian populations, consistent with the rapid evolution of PRDM9.

**Table 1.**
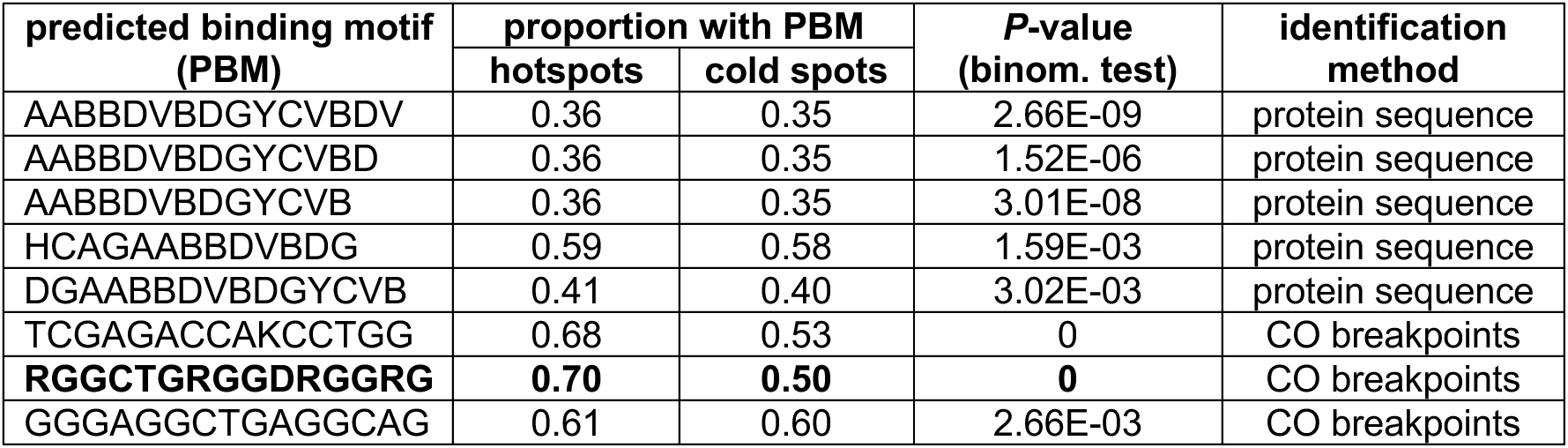
Predicted PRDM9 binding motifs (PBMs) with significant enrichment in recombination hotspots compared to sequence-matched cold spots in the Indian population (note: no significant enrichment was observed in the Chinese population). PBMs were predicted based on the position weight matrix predicted from the published PRDM9 protein sequence or the crossover breakpoints observed in pedigreed individuals. The best predicted PBM is highlighted in bold.

Finally, we used AlphaFold3’s machine learning guided model to predict the interaction between the best predicted PRDM9 binding motif — the 15 bp degenerate sequence RGGCTGAGGAGGRGG — and the PRDM9 sequence of rhesus macaques. In agreement with the strong enrichment observed within hotspots in the Indian population (with the motif being present in 70.5% of hotspots compared to 49.8% of sequence-matched cold spots; binomial test *P*-value = 0; Table 1), AlphaFold3 predicted a stable protein-DNA complex with high confidence (ipTM = 0.87; with 1.0 being the maximum value of the ipTM performance metric; Figure 4D).

### Correlations with genomic features

Previous studies in a variety of species have demonstrated that recombination rates tend to correlate with several genomic features, including nucleotide diversity, divergence, GC-content, repeat-content, and gene-content (e.g., Jensen-Seaman et al. 2004; Stevison and Noor 2010; Pfeifer and Jensen 2016; Stevison et al. 2016; Pfeifer 2020a; and see the review by Coop and Przeworski 2007). Consistent with this, we observed multiple significant correlations, with the strongest correlations generally observed at the broad (1Mb) scale (Figure 5). In both populations, recombination rates were most strongly and significantly correlated with GC-content, with the strength of the correlations ranging between 0.29 at the 10kb-scale and 0.51 at the 1Mb-scale. This pattern is likely driven by GC-biased gene conversion, which favors G and C alleles over A and T alleles when correcting DNA mismatches arising from recombination (see the reviews by Marais 2003; Duret and Galtier 2009; Galtier et al. 2009). In agreement with prior studies in primates (Pfeifer and Jensen 2016; Stevison et al. 2016; Soni et al 2025a,b), divergence (particularly to *M. fascicularis*) and nucleotide diversity were both significantly positively correlated with recombination rates, with correlation rates in the former ranging from 0.19 at the 10kb-scale to 0.40 at the 1Mb-scale and in the latter ranging from 0.08 at the 10kb-scale to 0.20 at the 1Mb-scale. A positive correlation between recombination rate and divergence is expected due to the mutagenicity of both crossover and non-crossover recombination (Halldorsson et al. 2019; Palsson et al. 2025). Additionally, nucleotide diversity is expected to correlate positively with recombination rates because of their interaction with natural selection (Cutter and Payseur 2013; Charlesworth and Jensen 2021). For example, the effects of purifying selection on linked sites (i.e., background selection; Charlesworth et al. 1993) is modulated by the local recombination rate. Consequently, regions of low recombination experience greater losses of neutral diversity, whereas regions of high recombination retain higher nucleotide diversity (Charlesworth et al. 1993). A similar pattern arises under selective sweeps: when a beneficial mutation fixes, linked variants may be carried to fixation as well (Maynard Smith and Haigh 1974). In low-recombination regions, this effect spans larger genomic segments, substantially reducing nucleotide diversity, while higher recombination restricts the impact to a narrower region, preserving more variation (Begun and Aquadro 1992). A less prominent positive correlation also exists with repeat-content, ranging from 0.01 at the 10kb-scale to 0.12 at the 1Mb-scale, as repetitive regions can act as hotspots for double-strand breaks that are subsequentially repaired by non-allelic homologous recombination (see the review by McVean 2010). Lastly, a negative correlation was observed with gene density, a correlation that is stronger at the fine (10kb) scale (−0.10) as PRDM9 directs recombination away from functional regions (Myers et al. 2005; Berg et al. 2010; Brick et al. 2012).

**Figure 5.**
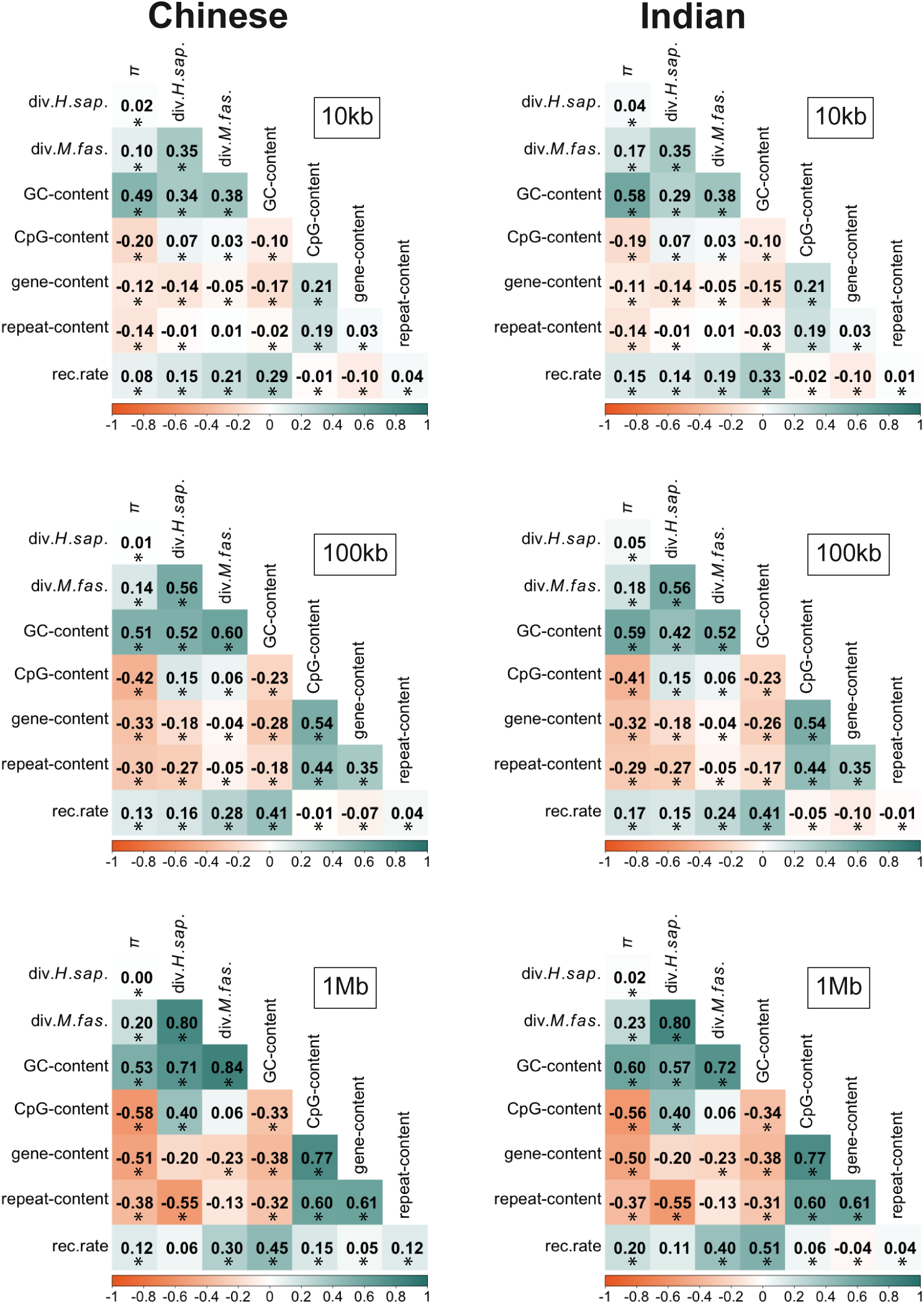
Genome-wide Spearman’s rank correlations between population-specific recombination rates inferred from the long-read data using LDhat and several genomic features, including nucleotide diversity, divergence from both *M. fascicularis* and *H. sapiens*, GC-content, CpG-content, gene-content, and repeat-content at the 10 kb, 100 kb, and 1 Mb scales. Positive correlations are shown in teal and negative correlations in orange, with the color intensity being proportional to the strength of the correlation. Significant correlations (*P*-value < 0.05) are marked with an asterisk.

## Conclusion

By generating both fine-scale mutation and recombination rate maps, we provide further insights into these fundamental evolutionary processes in a species central to biomedical and translational research, the rhesus macaque (*Macaca mulatta*). Specifically, we identified two possible ranges of fine-scale mutation rates based on divergence from one closely related (*M. fascicularis*) and one more distantly related (*H. sapiens*) primate species. Previously estimated pedigree-based rates more closely align with the fine-scale mutation rates inferred based on the divergence from *M. fascicularis*, suggesting that the long-time genome-wide average mutation rate of rhesus macaques likely falls between 0.32 × 10^-8^ and 0.62 × 10^-8^ /bp/gen. Accordingly, our results support a more recent split time of 2.7 mya between *M. mulatta* and *M. fascicularis*, and generation times between 9 and 11 years in the species. In addition, leveraging both short-(Illumina) and long-(PacBio HiFi) read sequencing data, we characterized the heterogeneity in recombination rates across the rhesus macaque genome demonstrating that long-read sequencing provides more accurate and higher-resolution estimates of recombination rates, improving the ability to detect fine-scale variation both within and between populations. Moreover, although broad-scale features of the recombination landscape are well-conserved between different rhesus macaque populations as well as with other primates, notable differences exist at finer scales, with the majority of recombination hotspots being private to a single population. Given the vital importance of population-specific estimates in population genomics, the results presented here will thus act as a foundation for future studies that require accurate mutation and recombination rate maps, including analyses of demography and selection as well as association and fine mapping studies of disease genes in populations of both Chinese and Indian descent housed at the U.S. Primate Research Centers and beyond.

## Acknowledgements

We would like to thank Sam Peterson and the team at the Oregon National Primate Research Center (ONPRC) for providing the rhesus macaque samples used in this study. DNA extraction was performed at the ONPRC (Beaverton, OR, USA), library preparation and PacBio HiFi sequencing were performed at the Arizona Genomics Institute at the University of Arizona (Tucson, AZ, USA). Computations were performed on the Sol supercomputer at Arizona State University (Jennewein et al. 2023).

## References

1000 Genomes Project Consortium. 2010. A map of human genome variation from population-scale sequencing. Nature. 467 (7319): 1061–1073. 10.1038/nature09534.

1000 Genomes Project Consortium. 2015. A global reference for human genetic variation. Nature. 526 (7571): 68–74. 10.1038/nature15393.

Abramson J, Adler J, Dunger J, Evans R, Green T, Pritzel A, Ronneberger O, Willmore L, Ballard AJ, Bambrick J, et al. 2024. Accurate structure prediction of biomolecular interactions with AlphaFold 3. Nature. 630 (8016): 493–500. 10.1038/s41586-024-07487-w.

Amarasinghe SL, Su S, Dong X, Zappia L, Ritchie ME, Gouil Q. 2020. Opportunities and challenges in long-read sequencing data analysis. Genome Biology. 21 (1): 30. 10.1186/s13059-020-1935-5.

Arbogast BS, Edwards SV, Wakeley J, Beerli P, Slowinski JB. 2002. Estimating divergence times from molecular data on phylogenetic and population genetic timescales. Annual Review of Ecology and Systematics. 33 (1): 707–740. 10.1146/annurev.ecolsys.33.010802.150500.

Armstrong J, Hickey G, Diekhans M, Fiddes IT, Novak AM, Deran A, Fang Q, Xie D, Feng S, Stiller J, et al. 2020. Progressive Cactus is a multiple-genome aligner for the thousand-genome era. Nature. 587 (7833): 246–251. 10.1038/s41586-020-2871-y.

Auton A, Fledel-Alon A, Pfeifer S, Venn O, Ségurel L, Street T, Leffler EM, Bowden R, Aneas I, Broxholme J, Humburg P, et al. 2012. A fine-scale chimpanzee genetic map from population sequencing. Science. 336 (6078): 193–198. 10.1126/science.1216872.

Auton A, McVean G. 2007. Recombination rate estimation in the presence of hotspots. Genome Research. 17 (8): 1219–1227. 10.1101/gr.6386707.

Bailey TL, Elkan C. 1994. Fitting a mixture model by expectation maximization to discover motifs in biopolymers. Proceedings of the International Conference on Intelligent Systems for Molecular Biology. 2: 28–36.

Bailey TL, Johnson J, Grant CE, Noble WS. 2015. The MEME suite. Nucleic Acids Research. 43 (W1): W39–W49. 10.1093/nar/gkv416.

Baudat F, Buard J, Grey C, Fledel-Alon A, Ober C, Przeworski M, Coop G, de Massy B. 2010. PRDM9 is a major determinant of meiotic recombination hotspots in humans and mice. Science. 327 (5967): 836–840. 10.1126/science.1183439.

Baumdicker F, Bisschop G, Goldstein D, Gower G, Ragsdale AP, Tsambos G, Zhu S, Eldon B, Ellerman EC, Galloway JG, et al. 2022. Efficient ancestry and mutation simulation with msprime 1.0. Genetics. 220 (3): iyab229. 10.1093/genetics/iyab229.

Begun DJ, Aquadro CF. 1992. Levels of naturally occurring DNA polymorphism correlate with recombination rates in *D. melanogaster*. Nature. 356 (6369): 519–520. 10.1038/356519a0.

Berg IL, Neumann R, Lam KW, Sarbajna S, Odenthal-Hesse L, May CA, Jeffreys AJ. 2010. PRDM9 variation strongly influences recombination hot-spot activity and meiotic instability in humans. Nature Genetics. 42 (10): 859–863. 10.1038/ng.658.

Bergeron LA, Besenbacher S, Bakker J, Zheng J, Li P, Pacheco G, Sinding MS, Kamilari M, Gilbert MTP, Schierup MH, et al. 2021. The germline mutational process in rhesus macaque and its implications for phylogenetic dating. GigaScience 10 (5): giab029. 10.1093/gigascience/giab029.

Bergeron LA, Besenbacher S, Zheng J, Li P, Bertelsen MF, Quintard B, Hoffman JI, Li Z, St Leger J, Shao C, et al. 2023. Evolution of the germline mutation rate across vertebrates. Nature. 615 (7951): 285–291. 10.1038/s41586-023-05752-y.

Besenbacher S, Hvilsom C, Marques-Bonet T, Mailund T, Schierup MH. 2019. Direct estimation of mutations in great apes reconciles phylogenetic dating. Nature Ecology and Evolution. 3 (2): 286–292. 10.1038/s41559-018-0778-x.

Bimber BN, Yan MY, Peterson SM, Ferguson B. 2019. mGAP: the macaque genotype and phenotype resource, a framework for accessing and interpreting macaque variant data, and identifying new models of human disease. BMC Genomics. 20 (1): 176. 10.1186/s12864-019-5559-7.

Brick K, Smagulova F, Khil P, Camerini-Otero RD, Petukhova GV. 2012. Genetic recombination is directed away from functional genomic elements in mice. Nature. 485 (7400): 642–645. 10.1038/nature11089.

Browne PD, Nielsen TK, Kot W, Aggerholm A, Gilbert MTP, Puetz L, Rasmussen M, Zervas A, Hansen LH. 2020. GC bias affects genomic and metagenomic reconstructions, underrepresenting GC-poor organisms. GigaScience 9 (2): giaa008. 10.1093/gigascience/giaa008.

Browning BL, Tian X, Zhou Y, Browning SR. 2021. Fast two-stage phasing of large-scale sequence data. The American Journal of Human Genetics. 108(10): 1880–1890. 10.1016/j.ajhg.2021.08.005.

Casper J, Speir ML, Raney BJ, Perez G, Nassar LR, Lee CM, Hinrichs AS, Gonzalez JN, Fischer C, Diekhans M, et al. 2026. The UCSC Genome Browser database: 2026 update. Nucleic Acids Research. 54(D1): D1331–D1335. 10.1093/nar/gkaf1250.

Champoux M, Higley JD, Suomi SJ. 1997. Behavioral and physiological characteristics of Indian and Chinese-Indian hybrid rhesus macaque infants. Developmental Psychobiology. 31 (1): 49–63.

Chang CC, Chow CC, Tellier LC, Vattikuti S, Purcell SM, Lee JJ. 2015. Second-generation PLINK: rising to the challenge of larger and richer datasets. GigaScience 4 (1): s13742-015-0047–0048. 10.1186/s13742-015-0047-8.

Charlesworth B, Jensen JD. 2021. Effects of selection at linked sites on patterns of genetic variability. Annual Review of Ecology, Evolution, and Systematics. 52 (1): 177–197. 10.1146/annurev-ecolsys-010621-044528.

Charlesworth B, Morgan MT, Charlesworth D. 1993. The effect of deleterious mutations on neutral molecular variation. Genetics. 134 (4): 1289–1303. 10.1093/genetics/134.4.1289.

Chintalapati M, Moorjani P. 2020. Evolution of the mutation rate across primates. Current Opinion in Genetics & Development. 62: 58–64. 10.1016/j.gde.2020.05.028.

Clarke MR, O’Neil JAS. 1999. Morphometric comparison of Chinese-origin and Indian-derived rhesus monkeys (*Macaca mulatta*). American Journal of Primatology. 47 (4): 335–346. 10.1002/(SICI)1098-2345(1999)47:4%253C335::AID-AJP5%253E3.0.CO;2-Y.

Cock PJA, Antao T, Chang JT, Chapman BA, Cox CJ, Dalke A, Friedberg I, Hamelryck T, Kauff F, Wilczynski B, et al. 2009. Biopython: freely available python tools for computational molecular biology and bioinformatics. Bioinformatics. 25 (11): 1422–1423. 10.1093/bioinformatics/btp163.

Conrad DF, Keebler JE, DePristo MA, Lindsay SJ, Zhang Y, Casals F, Idaghdour Y, Hartl CL, Torroja C, Garimella KV, et al. 2011. Variation in genome-wide mutation rates within and between human families. Nature Genetics. 43 (7): 712–714. 10.1038/ng.862.

Coop G, Przeworski M. 2007. An evolutionary view of human recombination. Nature Reviews. Genetics. 8 (1): 23–34. 10.1038/nrg1947.

Cooper EB, Brent LJ, Snyder-Mackler N, Singh M, Sengupta A, Khatiwada S, Malaivijitnond S, Qi Hai Z, Higham JP. 2022. The rhesus macaque as a success story of the Anthropocene. eLife. 11: e78169. 10.7554/eLife.78169.

Crow JF. 2000. The origins, patterns and implications of human spontaneous mutation. Nature Reviews. Genetics. 1 (1): 40–47. 10.1038/35049558.

Cutter AD, Payseur BA. 2013. Genomic signatures of selection at linked sites: unifying the disparity among species. Nature Reviews Genetics. 14 (4): 262–274. 10.1038/nrg3425.

Danecek P, Auton A, Abecasis G, Albers CA, Banks E, DePristo MA, Handsaker RE, Lunter G, Marth GT, Sherry ST, et al. 2011. The variant call format and VCFtools. Bioinformatics. 27 (15): 2156–2158. 10.1093/bioinformatics/btr330.

Danecek P, Bonfield JK, Liddle J, Marshall J, Ohan V, Pollard MO, Whitwham A, Keane T, McCarthy SA, Davies RM, et al. 2021. Twelve years of SAMtools and BCFtools. GigaScience. 10 (2): giab008. 10.1093/gigascience/giab008.

Dapper AL, Payseur BA. 2018. Effects of demographic history on the detection of recombination hotspots from linkage disequilibrium. Molecular Biology and Evolution. 35 (2): 335–353. 10.1093/molbev/msx272.

De Coster W, Weissensteiner MH, Sedlazeck FJ. 2021. Towards population-scale long-read sequencing. Nature Reviews Genetics. 22 (9): 572–587. 10.1038/s41576-021-00367-3.

Disotell TR, Tosi AJ. 2007. The monkey’s perspective. Genome Biology. 8 (9): 226. 10.1186/gb-2007-8-9-226.

Doxiadis GG, Otting N, de Groot NG, Bontrop RE. 2001. Differential evolutionary MHC class II strategies in humans and rhesus macaques: relevance for biomedical studies. Immunological Reviews. 183: 76–85. 10.1034/j.1600-065x.2001.1830106.x.

Doxiadis GG, Otting N, de Groot NG, de Groot N, Rouweler AJ, Noort R, Verschoor EJ, Bontjer I, Bontrop RE. 2003. Evolutionary stability of MHC class II haplotypes in diverse rhesus macaque populations. Immunogenetics. 55 (8): 540–551. 10.1007/s00251-003-0590-9.

Duret L, Galtier N. 2009. Biased gene conversion and the evolution of mammalian genomic landscapes. Annual Review of Genomics and Human Genetics. 10 (1): 285–311. 10.1146/annurev-genom-082908-150001.

Dutheil JY. 2024. On the estimation of genome-average recombination rates. Genetics. 227 (2): iyae051. 10.1093/genetics/iyae051.

Eggertsson HP, Jonsson H, Kristmundsdottir S, Hjartarson E, Kehr B, Masson G, Zink F, Hjorleifsson KE, Jonasdottir A, Jonasdottir A, et al. 2017. Graphtyper enables population-scale genotyping using pangenome graphs. Nature Genetics. 49 (11): 1654–1660. 10.1038/ng.3964.

Eggertsson HP, Kristmundsdottir S, Beyter D, Jonsson H, Skuladottir A, Hardarson MT, Gudbjartsson DF, Stefansson K, Halldorsson BV, Melsted P. 2019. GraphTyper2 enables population-scale genotyping of structural variation using pangenome graphs. Nature Communications. 10 (1): 5402. 10.1038/s41467-019-13341-9.

Espinosa E, Bautista R, Larrosa R, Plata O. 2024. Advancements in long-read genome sequencing technologies and algorithms. Genomics. 116 (3): 110842. 10.1016/j.ygeno.2024.110842.

Felsenstein J. 1974. The evolutionary advantage of recombination. Genetics. 78 (2): 737–756. 10.1093/genetics/78.2.737.

Gagliardi C, Liukkonen JR, Phillippi-Falkenstein KM, Harrison RM, Kubisch HM. 2007. Age as a determinant of reproductive success among captive female rhesus macaques (*Macaca mulatta*). Reproduction. 133 (4): 819–826. 10.1530/REP-06-0323.

Galtier N, Duret L, Glémin S, Ranwez V. 2009. GC-biased gene conversion promotes the fixation of deleterious amino acid changes in primates. Trends in Genetics. 25 (1): 1–5. 10.1016/j.tig.2008.10.011.

Goodman M, Porter CA, Czelusniak J, Page SL, Schneider H, Shoshani J, Gunnell G, Groves CP. 1998. Toward a phylogenetic classification of primates based on DNA evidence complemented by fossil evidence. Molecular Phylogenetics and Evolution. 9 (3): 585–598. 10.1006/mpev.1998.0495.

Grant CE, Bailey TL, Noble WS. 2011. FIMO: scanning for occurrences of a given motif. Bioinformatics. 27 (7): 1017–1018. 10.1093/bioinformatics/btr064.

Groves C. 2001. Primate taxonomy. Washington Smithsonian Institution Press. p. 229–232.

Halldorsson BV, Hardarson MT, Kehr B, Styrkarsdottir U, Gylfason A, Thorleifsson G, Zink F, Jonasdottir A, Jonasdottir A, Sulem P, et al. 2016. The rate of meiotic gene conversion varies by sex and age. Nature Genetics. 48 (11): 1377–1384. 10.1038/ng.3669.

Halldorsson BV, Palsson G, Stefansson OA, Jonsson H, Hardarson MT, Eggertsson HP, Gunnarsson B, Oddsson A, Halldorsson GH, Zink F, et al. 2019. Characterizing mutagenic effects of recombination through a sequence-level genetic map. Science. 363 (6425): eaau1043. 10.1126/science.aau1043.

Heenkenda EJ, Versoza CJ, Terbot JW, Soni V, Spatola GJ, Pfeifer SP, Jensen JD. 2026. Inferring the demographic history of Chinese and Indian rhesus macaque (Macaca mulatta) populations from PacBio HiFi long-read sequencing data. bioRxiv, preprint.

Hernandez RD, Hubisz MJ, Wheeler DA, Smith DG, Ferguson B, Rogers J, Nazareth L, Indap A, Bourquin T, McPherson J, et al. 2007. Demographic histories and patterns of linkage disequilibrium in Chinese and Indian rhesus macaques. Science. 316 (5822): 240–243. 10.1126/science.1140462.

Hill WG, Robertson A. 1966. The effect of linkage on limits to artificial selection. Genetical Research. 8 (3): 269–294.

Hinch AG, Tandon A, Patterson N, Song Y, Rohland N, Palmer CD, Chen GK, Wang K, Buxbaum SG, Akylbekova EL, et al. 2011. The landscape of recombination in African Americans. Nature. 476 (7359): 170–175. 10.1038/nature10336.

Hoge C, De Manuel M, Mahgoub M, Okami N, Fuller Z, Banerjee S, Baker Z, McNulty M, Andolfatto P, Macfarlan TS, et al. 2024. Patterns of recombination in snakes reveal a tug-of-war between PRDM9 and promoter-like features. Science. 383(6685): eadj7026. 10.1126/science.adj7026.

International HapMap Consortium. 2005. A haplotype map of the human genome. Nature. 437 (7063): 1299–1320. 10.1038/nature04226.

International HapMap Consortium. 2007. A second generation human haplotype map of over 3.1 million SNPs. Nature. 449 (7164): 851–861. 10.1038/nature06258.

Jennewein D, Lee J, Kurtz C, Dizon W, Shaeffer I, Chapman A, Chiquete A, Burks J, Carlson A, Mason N, et al. 2023. The Sol Supercomputer at Arizona State University. In: Practice and experience in advanced research computing. ACM: Portland, OR, USA. pp.296–301. 10.1145/3569951.3597573.

Jensen JD. 2023. Population genetic concerns related to the interpretation of empirical outliers and the neglect of common evolutionary processes. Heredity 130 (3): 109–110. 10.1038/s41437-022-00575-5.

Jensen-Seaman MI, Furey TS, Payseur BA, Lu Y, Roskin KM, Chen CF, Thomas MA, Haussler D, Jacob HJ. 2004. Comparative recombination rates in the rat, mouse, and human genomes. Genome Research. 14 (4): 528–538. 10.1101/gr.1970304.

Johnston HR, Cutler DJ. 2012. Population demographic history can cause the appearance of recombination hotspots. The American Journal of Human Genetics. 90 (5): 774–783. 10.1016/j.ajhg.2012.03.011.

Johnston SE. 2024. Understanding the genetic basis of variation in meiotic recombination: past, present, and future. Molecular Biology and Evolution. 41 (7): msae112. 10.1093/molbev/msae112.

Johri P, Aquadro CF, Beaumont M, Charlesworth B, Excoffier L, Eyre-Walker A, Keightley PD, Lynch M, McVean G, Payseur BA, et al. 2022a. Recommendations for improving statistical inference in population genomics. PLoS Biology. 20 (5): e3001669. 10.1371/journal.pbio.3001669.

Johri P, Eyre-Walker A, Gutenkunst RN, Lohmueller KE, Jensen JD. 2022b. On the prospect of achieving accurate joint estimation of selection with population history. Genome Biology and Evolution. 14 (7): evac088. 10.1093/gbe/evac088.

Kamm JA, Spence JP, Chan J, Song YS. 2016. Two-locus likelihoods under variable population size and fine-scale recombination rate estimation. Genetics. 203 (3): 1381–1399. 10.1534/genetics.115.184820.

Kassambara A. 2023. rstatix: pipe-friendly framework for basic statistical tests. R package version 0.7.2, https://rpkgs.datanovia.com/rstatix/.

Kimura M. 1968. Evolutionary rate at the molecular level. Nature. 217 (5129): 624–626. 10.1038/217624a0.

Korhonen J, Martinmäki P, Pizzi C, Rastas P, Ukkonen E. 2009. MOODS: fast search for position weight matrix matches in DNA sequences. Bioinformatics. 25(23): 3181–3182. 10.1093/bioinformatics/btp554.

Korunes KL, Samuk K. 2021. pixy: unbiased estimation of nucleotide diversity and divergence in the presence of missing data. Molecular Ecology Resources. 21 (4): 1359–1368. 10.1111/1755-0998.13326.

Kuderna LFK, Ulirsch JC, Rashid S, Ameen M, Sundaram L, Hickey G, Cox AJ, Gao H, Kumar A, Aguet F, et al. 2024. Identification of constrained sequence elements across 239 primate genomes. Nature. 625 (7996): 735–742. 10.1038/s41586-023-06798-8.

Lawrence M, Huber W, Pagès H, Aboyoun P, Carlson M, Gentleman R, Morgan MT, Carey VJ. 2013. Software for computing and annotating genomic ranges. PLoS Computational Biology. 9(8): e1003118. 10.1371/journal.pcbi.1003118.

Li H. 2013. Aligning sequence reads, clone sequences and assembly contigs with BWA-MEM. Preprint, arXiv. 10.48550/arXiv.1303.3997.

Li H. 2018. Minimap2: pairwise alignment for nucleotide sequences. Bioinformatics. 34 (18): 3094–3100. 10.1093/bioinformatics/bty191.

Manichaikul A, Mychaleckyj JC, Rich SS, Daly K, Sale M, Chen WM. 2010. Robust relationship inference in genome-wide association studies. Bioinformatics. 26 (22): 2867–2873. 10.1093/bioinformatics/btq559.

Marais G. 2003. Biased gene conversion: implications for genome and sex evolution. Trends in Genetics. 19 (6): 330–338. 10.1016/S0168-9525(03)00116-1.

Martin M, Patterson M, Garg S, O Fischer S, Pisanti N, Marschall T. 2016. WhatsHap: fast and accurate read-based phasing. Preprint, BioRxiv. 10.1101/085050.

Maruki T, Versoza CJ, Jensen JD, Pfeifer SP. 2026 Evolutionary genomics based on PacBio HiFi long-read sequencing data reveals the importance of structural variants in shaping population-specific differences between Chinese and Indian rhesus macaques (*Macaca mulatta*). bioRxiv, preprint.

Maynard Smith J, Haigh J. 1974. The hitch-hiking effect of a favourable gene. Genetical Research. 23 (1): 23–35.

McVean G. 2010. What drives recombination hotspots to repeat DNA in humans? Philosophical Transactions of the Royal Society of London, Series B, Biological Sciences. 365 (1544): 1213–1218. 10.1098/rstb.2009.0299.

Muller HJ. 1964. The relation of recombination to mutational advance. Mutation Research 1 (1): 2–9. 10.1016/0027-5107(64)90047-8.

Myers S, Bottolo L, Freeman C, McVean G, Donnelly P. 2005. A fine-scale map of recombination rates and hotspots across the human genome. Science. 310 (5746): 321–324. 10.1126/science.1117196.

Myers S, Bowden R, Tumian A, Bontrop RE, Freeman C, MacFie TS, McVean G, Donnelly P. 2010. Drive against hotspot motifs in primates implicates the *PRDM9* gene in meiotic recombination. Science. 327 (5967): 876–879. 10.1126/science.1182363.

Myers S, Freeman C, Auton A, Donnelly P, McVean G. 2008. A common sequence motif associated with recombination hot spots and genome instability in humans. Nature Genetics. 40 (9): 1124–1129. 10.1038/ng.213.

Nicolaisen LE, Desai MM. 2013. Distortions in genealogies due to purifying selection and recombination. Genetics. 195 (1): 221–230. 10.1534/genetics.113.152983.

O’Connell KA, Yosufzai ZB, Campbell RA, Lobb CJ, Engelken HT, Gorrell LM, Carlson TB, Catana JJ, Mikdadi D, Bonazzi VR, et al. 2023. Accelerating genomic workflows using NVIDIA Parabricks. BMC Bioinformatics. 24(1): 221. 10.1186/s12859-023-05292-2.

Paigen K, Petkov PM. 2018. PRDM9 and its role in genetic recombination. Trends in Genetics. 34 (4): 291–300. 10.1016/j.tig.2017.12.017.

Palsson G, Hardarson MT, Jonsson H, Steinthorsdottir V, Stefansson OA, Eggertsson HP, Gudjonsson SA, Olason PI, et al. 2025. Complete human recombination maps. Nature. 639 (8055): 700–707. 10.1038/s41586-024-08450-5.

Parvanov ED, Petkov PM, Paigen K. 2010. PRDM9 controls activation of mammalian recombination hotspots. Science. 327 (5967): 835. 10.1126/science.1181495.

Persikov AV, Osada R, Singh M. 2009. Predicting DNA recognition by Cys2His2 zinc finger proteins. Bioinformatics. 25(1): 22–29. 10.1093/bioinformatics/btn580.

Pfeifer SP. 2017. From next-generation resequencing reads to a high-quality variant data set. Heredity (Edinb). 118 (2): 111–124. 10.1038/hdy.2016.102.

Pfeifer SP. 2020a. A fine-scale genetic map for vervet monkeys. Molecular Biology and Evolution. 37 (7): 1855–1865. 10.1093/molbev/msaa079.

Pfeifer SP. 2020b. Spontaneous mutation rates. In The Molecular Evolutionary Clock, edited by Simon Y. W. Ho. Springer International Publishing. 10.1007/978-3-030-60181-2_3.

Pfeifer SP, Jensen JD. 2016. The impact of linked selection in chimpanzees: a comparative study. Genome Biology and Evolution. 8 (10): 3202–3208. 10.1093/gbe/evw240.

Poplin R, Chang PC, Alexander D, Schwartz S, Colthurst T, Ku A, Newburger D, Dijamco J, Nguyen N, Afshar PT, et al. 2018. A universal SNP and small-indel variant caller using deep neural networks. Nature Biotechnology. 36 (10): 983–987. 10.1038/nbt.4235.

Porsborg PS, Charmouh AP, Singh VK, Winge SB, Hvilsom C, Oroperv C, Hansen LT, Berner JA, Pelizzola M, Laurentino S, et al. 2025. Long-read sequencing of primate testis and human sperm allows identification of recombination events in individuals. Nature Communications. 16 (1): 10337. 10.1038/s41467-025-65248-3.

Pratto F, Brick K, Khil P, Smagulova F, Petukhova GV, Camerini-Otero DR. 2014. Recombination initiation maps of individual human genomes. Science. 346 (6211): 1256442. 10.1126/science.1256442.

Quinlan AR. 2014. BEDTools: the Swiss-army tool for genome feature analysis. Current Protocols in Bioinformatics. 47: 11.12.1–34. 10.1002/0471250953.bi1112s47.

R Core Team. 2025. R: a language and environment for statistical computing. R Foundation for Statistical Computing. Vienna, Austria. https://www.R-project.org/.

Rice P, Longden I, Bleasby A. 2000. EMBOSS: the European molecular biology open software suite. Trends in Genetics. 16 (6): 276–277. 10.1016/S0168-9525(00)02024-2.

Rogers J. 2022. Genomic resources for rhesus macaques (*Macaca mulatta*). Mammalian Genome. 33 (1): 91–99. 10.1007/s00335-021-09922-z.

Rogers J, Garcia R, Shelledy W, Kaplan J, Arya A, Johnson Z, Bergstrom M, Novakowski L, Nair P, Vinson A, et al. 2006. An initial genetic linkage map of the rhesus macaque (*Macaca mulatta*) genome using human microsatellite loci. Genomics. 87 (1): 30–38. 10.1016/j.ygeno.2005.10.004.

Samuk K, Noor MAF. 2022. Gene flow biases population genetic inference of recombination rate. G3 (Bethesda) 12 (11): jkac236. 10.1093/g3journal/jkac236.

Schwartz JJ, Roach DJ, Thomas JH, Shendure J. 2014. Primate evolution of the recombination regulator PRDM9. Nature Communications. 5 (1): 4370. 10.1038/ncomms5370.

Serre D, Nadon R, Hudson TJ. 2005. Large-scale recombination rate patterns are conserved among human populations. Genome Research. 15 (11): 1547–1552. 10.1101/gr.4211905.

Skipper M. 2007. Rhesus macaque joins the club. Nature Reviews Genetics. 8 (6): 410–411. 10.1038/nrg2121.

Slatko BE, Gardner AF, Ausubel FM. 2018. Overview of next-generation sequencing technologies. Current Protocols in Molecular Biology. 122 (1): e59. 10.1002/cpmb.59.

Soni V, Pfeifer SP, Jensen JD. 2024. The effects of mutation and recombination rate heterogeneity on the inference of demography and the distribution of fitness effects. Genome Biology and Evolution. 16 (2): evae004. 10.1093/gbe/evae004.

Soni V, Versoza CJ, Jensen JD, Pfeifer SP. 2025b. Inferring the landscapes of mutation and recombination in the common marmoset (*Callithrix jacchus*) in the presence of twinning and hematopoietic chimerism. Preprint, BioRxiv. 10.1101/2025.07.01.662565.

Soni V, Versoza CJ, Terbot JW, Jensen JD, Pfeifer SP. 2025a. Inferring fine-scale mutation and recombination rate maps in aye-ayes (*Daubentonia madagascariensis*). Ecology and Evolution. 15 (11): e72314. 10.1002/ece3.72314.

Soni V, Versoza CJ, Terbot JW, Spatola GJ, Bales KL, Jensen JD, Pfeifer SP. 2026. Inferring fine-scale rates of mutation and recombination in the coppery titi monkey (*Plecturocebus cupreus*). Preprint, BioRxiv. 10.64898/2026.01.13.699361.

Soni V, Versoza CJ, Vallender EJ, Jensen JD, Pfeifer SP. 2025c. Accounting for chimerism in demographic inference: reconstructing the history of common marmosets (*Callithrix jacchus*) from high-quality, whole-genome, population-level data. Molecular Biology and Evolution. 42 (6): msaf119. 10.1093/molbev/msaf119.

Spence JP, Song YS. 2019. Inference and analysis of population-specific fine-scale recombination maps across 26 diverse human populations. Science Advances. 5 (10): eaaw9206. 10.1126/sciadv.aaw9206.

Stapley J, Feulner PGD, Johnston SE, Santure AW, Smadja CM. 2017. Variation in recombination frequency and distribution across eukaryotes: patterns and processes. Philosophical Transactions of the Royal Society B: Biological Sciences. 372 (1736): 20160455. 10.1098/rstb.2016.0455.

Stevison LS, Noor MAF. 2010. Genetic and evolutionary correlates of fine-scale recombination rate variation in *Drosophila persimilis*. Journal of Molecular Evolution. 71 (5–6): 332–345. 10.1007/s00239-010-9388-1.

Stevison LS, Woerner AW, Kidd JM, Kelley JL, Veeramah KR, McManus KF; Great Ape Genome Project; Bustamante CD, Hammer MF, Wall JD. 2016. The time scale of recombination rate evolution in great apes. Molecular Biology and Evolution. 33 (4): 928–945. 10.1093/molbev/msv331.

Tan X, Qi J, Liu Z, Fan P, Liu G, Zhang L, Shen Y, Li J, Roos C, Zhou X, et al. 2023. Phylogenomics reveals high levels of incomplete lineage sorting at the ancestral nodes of the macaque radiation. Molecular Biology and Evolution. 40 (11): msad229. 10.1093/molbev/msad229.

Terbot JW, Calahorra-Oliart A, Versoza CJ, Shah D, Soni V, Pfeifer SP, Jensen JD. 2025a. Re-evaluating the demographic history of, and inferring the fine-scale recombination landscape for, wild Chinese rhesus macaques (*Macaca mulatta*). American Journal of Primatology. 87 (11): e70088. 10.1002/ajp.70088.

Terbot JW, Soni V, Versoza CJ, Pfeifer SP, Jensen JD. 2025b. Inferring the demographic history of aye-ayes (*Daubentonia madagascariensis*) from high-quality, whole-genome, population-level data. Genome Biology and Evolution. 17 (1): evae281. 10.1093/gbe/evae281.

Trichel AM, Rajakumar PA, Murphey-Corb M. 2002. Species-specific variation in SIV disease progression between Chinese and Indian subspecies of rhesus macaque. Journal of Medical Primatology. 31 (4-5): 171–178. 10.1034/j.1600-0684.2002.02003.x.

van der Auwera G, O’Connor BD. 2020. Genomics in the cloud: using Docker, GATK, and WDL in Terra. (First edition). O’Reilly Media.

Venn O, Turner I, Mathieson I, de Groot N, Bontrop R, McVean G. 2014. Strong male bias drives germline mutation in chimpanzees. Science. 344 (6189): 1272–1275. 10.1126/science.344.6189.1272.

Versoza CJ, Lloret-Villas A, Jensen JD, Pfeifer SP. 2025. A pedigree-based map of crossovers and noncrossovers in aye-ayes (*Daubentonia madagascariensis*). Genome Biology and Evolution. 17 (5): evaf072. 10.1093/gbe/evaf072.

Versoza CJ, Weiss A, Johal R, La Rosa B, Jensen JD, Pfeifer SP. 2024. Novel insights into the landscape of crossover and noncrossover events in rhesus macaques (*Macaca mulatta*). Genome Biology and Evolution. 16 (1): evad223. 10.1093/gbe/evad223.

Viray J, Rolfs B, Smith DG. 2001. Comparison of the frequencies of major histocompatibility (MHC) class-II DQA1 and DQB1 alleles in Indian and Chinese rhesus macaques (*Macaca mulatta*). Comparative Medicine. 51 (6): 555–561.

Wall JD, Robinson JA, Cox LA. 2022. High-resolution estimates of crossover and noncrossover recombination from a captive baboon colony. Genome Biology and Evolution. 14 (4): evac040. 10.1093/gbe/evac040.

Wang RJ, Thomas GWC, Raveendran M, Harris RA, Doddapaneni H, Muzny DM, Capitanio JP, Radivojac P, Rogers J, Hahn MW. 2020. Paternal age in rhesus macaques is positively associated with germline mutation accumulation but not with measures of offspring sociability. Genome Research. 30 (6): 826–834. 10.1101/gr.255174.119.

Warren WC, Harris RA, Haukness M, Fiddes IT, Murali SC, Fernandes J, Dishuck PC, Storer JM, Raveendran M, Hillier LW, et al. (2020). Sequence diversity analyses of an improved rhesus macaque genome enhance its biomedical utility. Science. 370(6523): eabc6617. 10.1126/science.abc6617.

Watterson GA. 1975. On the number of segregating sites in genetical models without recombination. Theoretical Population Biology. 7(2): 256–276. 10.1016/0040-5809(75)90020-9.

Williams AL, Genovese G, Dyer T, Altemose N, Truax K, Jun G, Patterson N, Myers SR, Curran JE, Duggirala R, et al. 2015. Non-crossover gene conversions show strong GC bias and unexpected clustering in humans. eLife 4: e04637. 10.7554/eLife.04637.

Xue C, Raveendran M, Harris RA, Fawcett GL, Liu X, White S, Dahdouli M, Rio Deiros D, Below JE, Salerno W, et al. 2016. The population genomics of rhesus macaques (*Macaca mulatta*) based on whole-genome sequences. Genome Research. 26 (12): 1651–1662. 10.1101/gr.204255.116.

Xue C, Rustagi N, Liu X, Raveendran M, Harris RA, Venkata MG, Rogers J, Yu F. 2020. Reduced meiotic recombination in rhesus macaques and the origin of the human recombination landscape. PLoS One 15 (8): e0236285. 10.1371/journal.pone.0236285.

Yun T, Li H, Chang PC, Lin MF, Carroll A, McLean CY. 2020. Accurate, scalable cohort variant calls using DeepVariant and GLnexus. Bioinformatics. 36 (24): 5582–5589. 10.1093/bioinformatics/btaa1081.

Zedek F, Bureš P, Elliott TL, Escudero M, Lucek K, Marques A. 2026. Chromosome size as a robust predictor of recombination rate: insights from holocentric and monocentric systems. Genetics. 232 (1): iyaf247. 10.1093/genetics/iyaf247.

Zhang S, Xu N, Fu L, Yang X, Ma K, Li Y, Yang Z, Li Z, Feng Y, Jiang X, et al. 2025. Integrated analysis of the complete sequence of a macaque genome. Nature. 640 (8059): 714–721. 10.1038/s41586-025-08596-w.

Zoonomia Consortium. 2020. A comparative genomics multitool for scientific discovery and conservation. Nature. 587 (7833): 240–245. 10.1038/s41586-020-2876-6.

