## Supplementary Materials for "Comparing fine-scale mutation and recombination landscapes in rhesus macaque (*Macaca mulatta*) populations of Chinese and Indian descent inferred from both short- and long-read sequencing data"

### Supplementary Tables

| population | method | dataset | demographic model | # sites with ≥ 100 $\rho$/kb |
|---|---|---|---|---|
| Chinese | LDhat | short-read | – | 6 |
| | | long-read | – | 215 |
| | pyrho | short-read | $\mu = 0.58 \times 10^{-8}$ | 0 |
| | | | $\mu = 1.49 \times 10^{-8}$ | 0 |
| | | long-read | $\mu = 0.58 \times 10^{-8}$ | 334 |
| | | | $\mu = 1.49 \times 10^{-8}$ | 238 |
| Indian | LDhat | short-read | – | 46 |
| | | long-read | – | 116 |
| | pyrho | short-read | $\mu = 0.58 \times 10^{-8}$ | 58,319 |
| | | | $\mu = 1.49 \times 10^{-8}$ | 68,095 |
| | | long-read | $\mu = 0.58 \times 10^{-8}$ | 50,178 |
| | | | $\mu = 1.49 \times 10^{-8}$ | 53,008 |

**Table S1.** The number of sites where population-scaled recombination rate estimates obtained using LDhat or pyrho were ≥ 100 $\rho$/kb. Recombination rates of these sites, as well as those of the surrounding 100 SNPs (i.e., 50 SNPs up- and down-stream), were masked by setting the rates to 0. Inference was based on both short- and long-read datasets for the Chinese and Indian rhesus macaque populations (see the sections entitled "Short-read data" and "Long-read data"). Estimates for the demography-aware estimator pyrho were based on the demographic histories recently inferred by Heenkenda et al. 2026 for these two populations, assuming mutation rates of $\mu = 0.58 \times 10^{-8}$ /bp/gen (based on the pedigree-based estimate of Wang et al. [2020]) or $\mu = 1.49 \times 10^{-8}$ /bp/gen (based on the maximum indirect estimate observed from patterns of *M. mulatta–H. sapiens* divergence in this study).

| window size | divergence (*H. sapiens*) | divergence (*M. fascicularis*) |
|---|---|---|
| 1 kb | 0.032420 (±) 0.010407 | 0.001468 (±) 0.001855 |
| 10 kb | 0.033043 (±) 0.005772 | 0.001498 (±) 0.000947 |
| 100 kb | 0.033536 (±) 0.004217 | 0.001531 (±) 0.000443 |
| 1 Mb | 0.033898 (±) 0.003486 | 0.001521 (±) 0.000259 |

**Table S2.** Mean neutral divergence from *H. sapiens* and *M. fascicularis* calculated in sliding windows of 1 kb, 10 kb, 100 kb, and 1 Mb across the rhesus macaque genome.

| mean mutation rate (/bp/gen) | window size (kb) | divergence time (mya) | generation time (years) | # windows |
|---|---|---|---|---|
| 4.89E-09 | 1 | 2.7 | 9 | 685,746 |
| 4.99E-09 | 10 | 2.7 | 9 | 46,870 |
| 5.10E-09 | 100 | 2.7 | 9 | 3,174 |
| 5.07E-09 | 1,000 | 2.7 | 9 | 171 |
| 3.15E-09 | 1 | 4.2 | 9 | 685,746 |
| 3.21E-09 | 10 | 4.2 | 9 | 46,870 |
| 3.28E-09 | 100 | 4.2 | 9 | 3,174 |
| 3.26E-09 | 1,000 | 4.2 | 9 | 171 |
| 5.98E-09 | 1 | 2.7 | 11 | 685,746 |
| 6.10E-09 | 10 | 2.7 | 11 | 46,870 |
| 6.24E-09 | 100 | 2.7 | 11 | 3,174 |
| 6.20E-09 | 1,000 | 2.7 | 11 | 171 |
| 3.85E-09 | 1 | 4.2 | 11 | 685,746 |
| 3.92E-09 | 10 | 4.2 | 11 | 46,870 |
| 4.01E-09 | 100 | 4.2 | 11 | 3,174 |
| 3.98E-09 | 1,000 | 4.2 | 11 | 171 |

**Table S3.** Summary of per-site per-generation mutation rate estimates averaged across 1 kb, 10 kb, 100 kb, and 1 Mb regions of the rhesus macaque genome based on generation times of 9 years and 11 years (Gagliardi et al. 2007; Xue et al. 2016) as well as divergence times for *M. mulatta–M. fascicularis* of 2.7 mya and 4.2 mya (Bergeron et al. 2021; Tan et al. 2023). # windows provides information about the number of accessible windows available at each window size.

| mean mutation rate (/bp/gen) | window size (kb) | divergence time (mya) | generation time (years) | # windows |
|---|---|---|---|---|
| 1.17E-08 | 1 | 25 | 9 | 590,755 |
| 1.19E-08 | 10 | 25 | 9 | 34,407 |
| 1.21E-08 | 100 | 25 | 9 | 2,111 |
| 1.22E-08 | 1,000 | 25 | 9 | 81 |
| 0.97E-08 | 1 | 30 | 9 | 590,755 |
| 0.99E-08 | 10 | 30 | 9 | 34,407 |
| 1.01E-08 | 100 | 30 | 9 | 2,111 |
| 1.02E-08 | 1,000 | 30 | 9 | 81 |
| 0.83E-08 | 1 | 35 | 9 | 590,755 |
| 0.85E-08 | 10 | 35 | 9 | 34,407 |
| 0.86E-08 | 100 | 35 | 9 | 2,111 |
| 0.87E-08 | 1,000 | 35 | 9 | 81 |
| 1.43E-08 | 1 | 25 | 11 | 590,755 |
| 1.45E-08 | 10 | 25 | 11 | 34,407 |
| 1.48E-08 | 100 | 25 | 11 | 2,111 |
| 1.49E-08 | 1,000 | 25 | 11 | 81 |
| 1.19E-08 | 1 | 30 | 11 | 590,755 |
| 1.21E-08 | 10 | 30 | 11 | 34,407 |
| 1.23E-08 | 100 | 30 | 11 | 2,111 |
| 1.24E-08 | 1,000 | 30 | 11 | 81 |
| 1.02E-08 | 1 | 35 | 11 | 590,755 |
| 1.04E-08 | 10 | 35 | 11 | 34,407 |
| 1.05E-08 | 100 | 35 | 11 | 2,111 |
| 1.07E-08 | 1,000 | 35 | 11 | 81 |

**Table S4.** Summary of per-site per-generation mutation rate estimates averaged across 1 kb, 10 kb, 100 kb, and 1 Mb regions of the rhesus macaque genome based on generation times of 9 years and 11 years (Gagliardi et al. 2007; Xue et al. 2016) as well as divergence times for *M. mulatta–H. sapiens* ranging from 25 mya to 35 mya (Goodman et al. 1998; Disotell and Tosi 2007; Skipper 2007; Chintalapati and Moorjani 2020). # windows provides information about the number of accessible windows available at each window size.

| population | mutation rate | dataset | method | mean $r$ (/bp/gen) | | | |
|---|---|---|---|---|---|---|---|
| | | | | 1 Mb | 100 kb | 10 kb | 1 kb |
| Chinese | $0.58 \times 10^{-8}$ | short-read | LDhat | $1.00 \times 10^{-8}$ | $0.94 \times 10^{-8}$ | $0.86 \times 10^{-8}$ | $0.89 \times 10^{-8}$ |
| | | | pyrho | $0.74 \times 10^{-8}$ | $0.72 \times 10^{-8}$ | $0.71 \times 10^{-8}$ | $0.71 \times 10^{-8}$ |
| | | long-read | LDhat | $0.66 \times 10^{-8}$ | $0.65 \times 10^{-8}$ | $0.64 \times 10^{-8}$ | $0.63 \times 10^{-8}$ |
| | | | pyrho | $0.63 \times 10^{-8}$ | $0.61 \times 10^{-8}$ | $0.60 \times 10^{-8}$ | $0.60 \times 10^{-8}$ |
| | $1.49 \times 10^{-8}$ | short-read | LDhat | $2.58 \times 10^{-8}$ | $2.40 \times 10^{-8}$ | $2.20 \times 10^{-8}$ | $2.29 \times 10^{-8}$ |
| | | | pyrho | $1.79 \times 10^{-8}$ | $1.73 \times 10^{-8}$ | $1.71 \times 10^{-8}$ | $1.71 \times 10^{-8}$ |
| | | long-read | LDhat | $1.70 \times 10^{-8}$ | $1.66 \times 10^{-8}$ | $1.63 \times 10^{-8}$ | $1.63 \times 10^{-8}$ |
| | | | pyrho | $1.52 \times 10^{-8}$ | $1.48 \times 10^{-8}$ | $1.46 \times 10^{-8}$ | $1.45 \times 10^{-8}$ |
| Indian | $0.58 \times 10^{-8}$ | short-read | LDhat | $0.24 \times 10^{-8}$ | $0.22 \times 10^{-8}$ | $0.21 \times 10^{-8}$ | $0.21 \times 10^{-8}$ |
| | | | pyrho | $1.40 \times 10^{-8}$ | $1.31 \times 10^{-8}$ | $1.24 \times 10^{-8}$ | $1.21 \times 10^{-8}$ |
| | | long-read | LDhat | $0.17 \times 10^{-8}$ | $0.16 \times 10^{-8}$ | $0.16 \times 10^{-8}$ | $0.16 \times 10^{-8}$ |
| | | | pyrho | $0.84 \times 10^{-8}$ | $0.81 \times 10^{-8}$ | $0.81 \times 10^{-8}$ | $0.84 \times 10^{-8}$ |
| | $1.49 \times 10^{-8}$ | short-read | LDhat | $0.60 \times 10^{-8}$ | $0.57 \times 10^{-8}$ | $0.54 \times 10^{-8}$ | $0.53 \times 10^{-8}$ |
| | | | pyrho | $3.17 \times 10^{-8}$ | $2.98 \times 10^{-8}$ | $2.83 \times 10^{-8}$ | $2.76 \times 10^{-8}$ |
| | | long-read | LDhat | $0.42 \times 10^{-8}$ | $0.41 \times 10^{-8}$ | $0.41 \times 10^{-8}$ | $0.40 \times 10^{-8}$ |
| | | | pyrho | $1.87 \times 10^{-8}$ | $1.82 \times 10^{-8}$ | $1.83 \times 10^{-8}$ | $1.92 \times 10^{-8}$ |

**Table S5.** Average per-site per-generation recombination rate estimates for rhesus macaque populations of Chinese and Indian descent as inferred from short-read and long-read data using LDhat and pyrho across 1 Mb, 100 kb, 10 kb, and 1 kb regions of the rhesus macaque genome. Population-scaled recombination rate estimates were converted to per-generation estimates using population-specific $N_e$ values based on mutation rates ($\mu$) of $0.58 \times 10^{-8}$ /bp/gen (i.e., the pedigree-based estimate obtained by Wang et al. [2020]) or $1.49 \times 10^{-8}$ /bp/gen (i.e., the maximum indirect estimate observed from patterns of *M. mulatta–H. sapiens* divergence in this study).

| population | mutation rate | mean $r$ (/bp/gen) | | | | # hotspots | |
|---|---|---|---|---|---|---|---|
| | | 1 Mb | 100 kb | 10 kb | 1 kb | 5× | 10× |
| Chinese | $0.58 \times 10^{-8}$ | $0.59 \times 10^{-8}$ | $0.57 \times 10^{-8}$ | $0.55 \times 10^{-8}$ | $0.54 \times 10^{-8}$ | 31,856 | 1,496 |
| | $1.49 \times 10^{-8}$ | $1.52 \times 10^{-8}$ | $1.48 \times 10^{-8}$ | $1.42 \times 10^{-8}$ | $1.38 \times 10^{-8}$ | | |
| Indian | $0.58 \times 10^{-8}$ | $0.16 \times 10^{-8}$ | $0.16 \times 10^{-8}$ | $0.15 \times 10^{-8}$ | $0.15 \times 10^{-8}$ | 86,158 | 21,597 |
| | $1.49 \times 10^{-8}$ | $0.42 \times 10^{-8}$ | $0.40 \times 10^{-8}$ | $0.39 \times 10^{-8}$ | $0.38 \times 10^{-8}$ | | |

**Table S6.** Average per-site per-generation recombination rate estimates for rhesus macaque populations of Chinese and Indian descent as inferred from long-read data including singletons using LDhat across 1 Mb, 100 kb, 10 kb, and 1 kb regions of the rhesus macaque genome. Population-scaled recombination rate estimates were converted to per-generation estimates using population-specific $N_e$ values based on mutation rates ($\mu$) of $0.58 \times 10^{-8}$ /bp/gen (i.e., the pedigree-based estimate obtained by Wang et al. [2020]) or $1.49 \times 10^{-8}$ /bp/gen (i.e., the maximum indirect estimate observed from patterns of *M. mulatta–H. sapiens* divergence in this study).

| population | mutation rate | dataset | method | scale | mean recombination rate (cM/Mb) |
|---|---|---|---|---|---|
| Chinese | $0.58 \times 10^{-8}$ | short-read | LDhat | 1 Mb | 0.909 (±) 0.111 |
| | | | pyrho | | 0.925 (±) 0.162 |
| | | long-read | LDhat | | 0.912 (±) 0.131 |
| | | | pyrho | | 0.924 (±) 0.171 |
| | $1.49 \times 10^{-8}$ | short-read | LDhat | | 0.909 (±) 0.111 |
| | | | pyrho | | 0.925 (±) 0.161 |
| | | long-read | LDhat | | 0.912 (±) 0.131 |
| | | | pyrho | | 0.924 (±) 0.172 |
| Indian | $0.58 \times 10^{-8}$ | short-read | LDhat | | 0.911 (±) 0.129 |
| | | | pyrho | | 0.904 (±) 0.125 |
| | | long-read | LDhat | | 0.913 (±) 0.136 |
| | | | pyrho | | 0.905 (±) 0.120 |
| | $1.49 \times 10^{-8}$ | short-read | LDhat | | 0.911 (±) 0.129 |
| | | | pyrho | | 0.904 (±) 0.131 |
| | | long-read | LDhat | | 0.913 (±) 0.136 |
| | | | pyrho | | 0.908 (±) 0.134 |

| population | mutation rate | dataset | method | scale | mean recombination rate (cM/Mb) |
|---|---|---|---|---|---|
| Chinese | $0.58 \times 10^{-8}$ | short-read | LDhat | 100 kb | 0.910 (±) 0.116 |
| | | | pyrho | | 0.925 (±) 0.163 |
| | | long-read | LDhat | | 0.912 (±) 0.133 |
| | | | pyrho | | 0.923 (±) 0.168 |
| | $1.49 \times 10^{-8}$ | short-read | LDhat | | 0.910 (±) 0.166 |
| | | | pyrho | | 0.925 (±) 0.162 |
| | | long-read | LDhat | | 0.912 (±) 0.133 |
| | | | pyrho | | 0.923 (±) 0.169 |
| Indian | $0.58 \times 10^{-8}$ | short-read | LDhat | | 0.912 (±) 0.131 |
| | | | pyrho | | 0.905 (±) 0.131 |
| | | long-read | LDhat | | 0.914 (±) 0.141 |
| | | | pyrho | | 0.906 (±) 0.129 |
| | $1.49 \times 10^{-8}$ | short-read | LDhat | | 0.912 (±) 0.131 |
| | | | pyrho | | 0.906 (±) 0.140 |
| | | long-read | LDhat | | 0.914 (±) 0.141 |
| | | | pyrho | | 0.910 (±) 0.144 |

| population | mutation rate | dataset | method | scale | mean recombination rate (cM/Mb) |
|---|---|---|---|---|---|
| Chinese | $0.58 \times 10^{-8}$ | short-read | LDhat | 10 kb | 0.912 (±) 0.124 |
| | | | pyrho | | 0.925 (±) 0.163 |
| | | long-read | LDhat | | 0.911 (±) 0.129 |
| | | | pyrho | | 0.923 (±) 0.165 |
| | $1.49 \times 10^{-8}$ | short-read | LDhat | | 0.912 (±) 0.124 |
| | | | pyrho | | 0.925 (±) 0.162 |
| | | long-read | LDhat | | 0.911 (±) 0.129 |
| | | | pyrho | | 0.923 (±) 0.166 |
| Indian | $0.58 \times 10^{-8}$ | short-read | LDhat | | 0.912 (±) 0.134 |
| | | | pyrho | | 0.906 (±) 0.132 |
| | | long-read | LDhat | | 0.914 (±) 0.143 |
| | | | pyrho | | 0.908 (±) 0.135 |
| | $1.49 \times 10^{-8}$ | short-read | LDhat | | 0.912 (±) 0.134 |
| | | | pyrho | | 0.907 (±) 0.140 |
| | | long-read | LDhat | | 0.914 (±) 0.143 |
| | | | pyrho | | 0.913 (±) 0.154 |

| population | mutation rate | dataset | method | scale | mean recombination rate (cM/Mb) |
|---|---|---|---|---|---|
| Chinese | $0.58 \times 10^{-8}$ | short-read | LDhat | 1 kb | 0.910 (±) 0.114 |
| | | | pyrho | | 0.925 (±) 0.163 |
| | | long-read | LDhat | | 0.911 (±) 0.131 |
| | | | pyrho | | 0.922 (±) 0.164 |
| | $1.49 \times 10^{-8}$ | short-read | LDhat | | 0.910 (±) 0.114 |
| | | | pyrho | | 0.925 (±) 0.162 |
| | | long-read | LDhat | | 0.911 (±) 0.131 |
| | | | pyrho | | 0.922 (±) 0.165 |
| Indian | $0.58 \times 10^{-8}$ | short-read | LDhat | | 0.912 (±) 0.131 |
| | | | pyrho | | 0.906 (±) 0.131 |
| | | long-read | LDhat | | 0.913 (±) 0.137 |
| | | | pyrho | | 0.910 (±) 0.139 |
| | $1.49 \times 10^{-8}$ | short-read | LDhat | | 0.912 (±) 0.131 |
| | | | pyrho | | 0.907 (±) 0.139 |
| | | long-read | LDhat | | 0.913 (±) 0.137 |
| | | | pyrho | | 0.915 (±) 0.158 |

**Table S7.** Average recombination rate estimates for rhesus macaque populations of Chinese and Indian descent as inferred from short-read and long-read data using LDhat and pyrho in 1 Mb, 100 kb, 10 kb, and 1 kb windows across the genome. Population-scaled recombination rate estimates were converted to per-generation estimates using population-specific $N_e$ values based on mutation rates ($\mu$) of $0.58 \times 10^{-8}$ /bp/gen (i.e., the pedigree-based estimate obtained by Wang et al. [2020]) or $1.49 \times 10^{-8}$ /bp/gen (i.e., the maximum indirect estimate observed from patterns of *M. mulatta*–*H. sapiens* divergence in this study) and re-scaled to the genetic map length previously observed from pedigree data (Versoza et al. 2024).

| population | heat | # hotspots | # cold spots | mean length (kb) | # shared hotspots |
|---|---|---|---|---|---|
| Chinese | > 5× | 34,101 | 33,976 | 2.82 | 9,032* |
| | > 10× | 5,136 | — | 2.31 | 721 |
| Indian | > 5× | 47,024 | 46,898 | 2.93 | 9,030 |
| | > 10× | 16,300 | — | 2.44 | 721 |

**Table S8.** Summary of recombination hotspots discovered in the Chinese and Indian rhesus macaque populations with a heat exceeding 5× or 10× the genomic background, the number of sequence-matched cold spots, the average length of recombination hotspots, and the number of hotspots shared between the two populations. * Note that one hotspot detected in the Chinese population overlapped with two hotspots in the Indian population and three hotspots detected in the Indian population overlapped with two hotspots in the Chinese population each, accounting for slight differences in the number of shared hotspots between the populations.

### Supplementary Figures

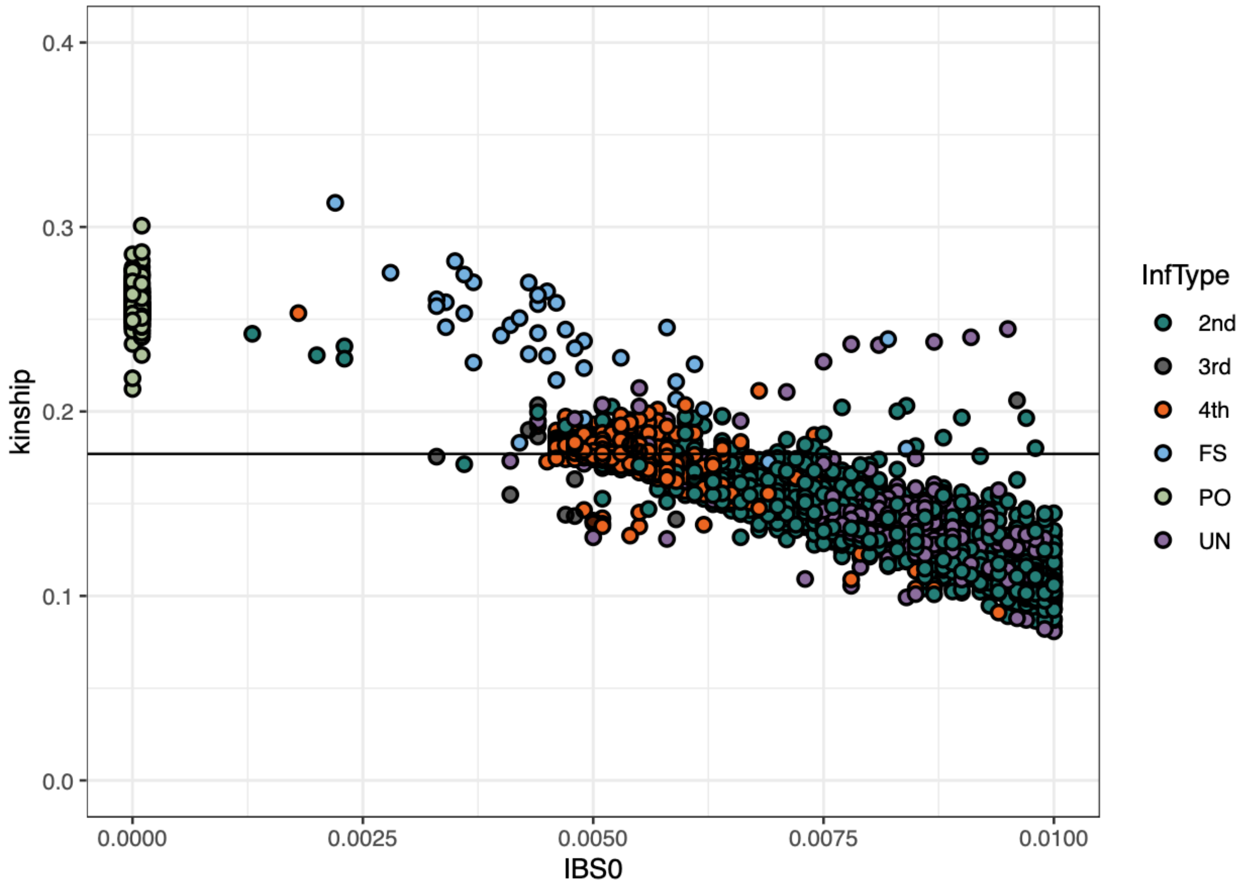


**Figure S1.** Scatterplot showing kinship coefficients and IBS0 (i.e., the total number of observations in which two discordant homozygotes are present) for each pairwise comparison among the 888 rhesus macaque individuals of Chinese and Indian descent included in the mGAP database (Bimber et al. 2019). Pairwise kinship coefficients greater than 0.177 (indicated by a solid black line) indicate a first-degree relationship (i.e., parent-offspring pairs or full siblings). InfType corresponds to relationship types inferred with KING (Manichaikul et al. 2010), with PO denoting parent-offspring pairs (shown in green), FS denoting full siblings (blue), 2nd denoting second-degree relationships (teal), 3rd denoting third-degree relationships (gray), 4th denoting fourth-degree relationships (orange), and UN denoting unrelated individuals (purple).

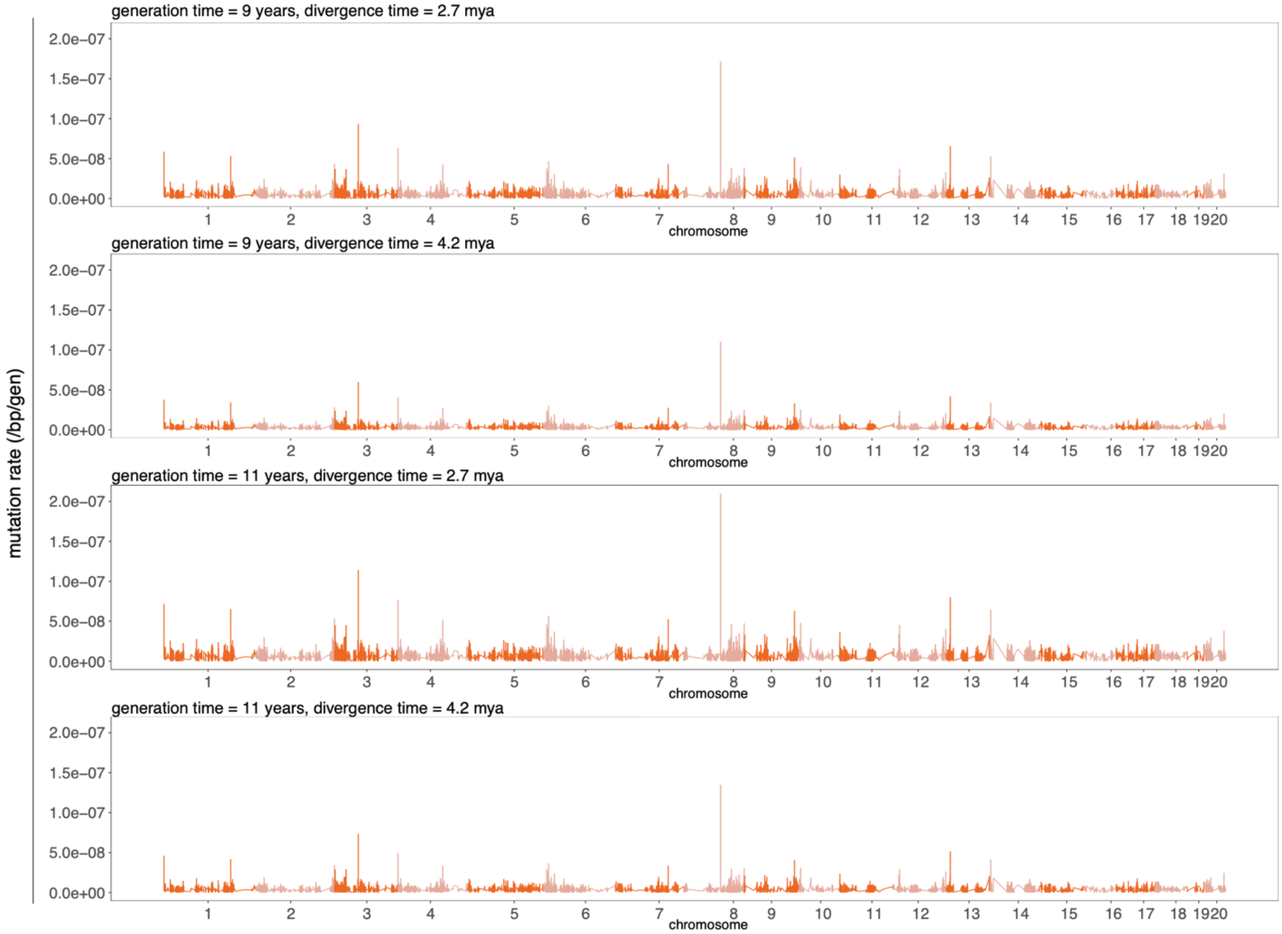


**Figure S2.** Fine-scale per-site per-generation mutation rate estimates observed in 10 kb windows with a step size of 5 kb based on patterns of neutral divergence between *M. mulatta* and *M. fascicularis*, assuming generation times of 9 years and 11 years (Gagliardi et al. 2007; Xue et al. 2016) as well as divergence times of 2.7 mya and 4.2 mya for *M. mulatta*–*M. fascicularis* (Bergeron et al. 2021; Tan et al. 2023). Individual chromosomes are shown in alternating shading.

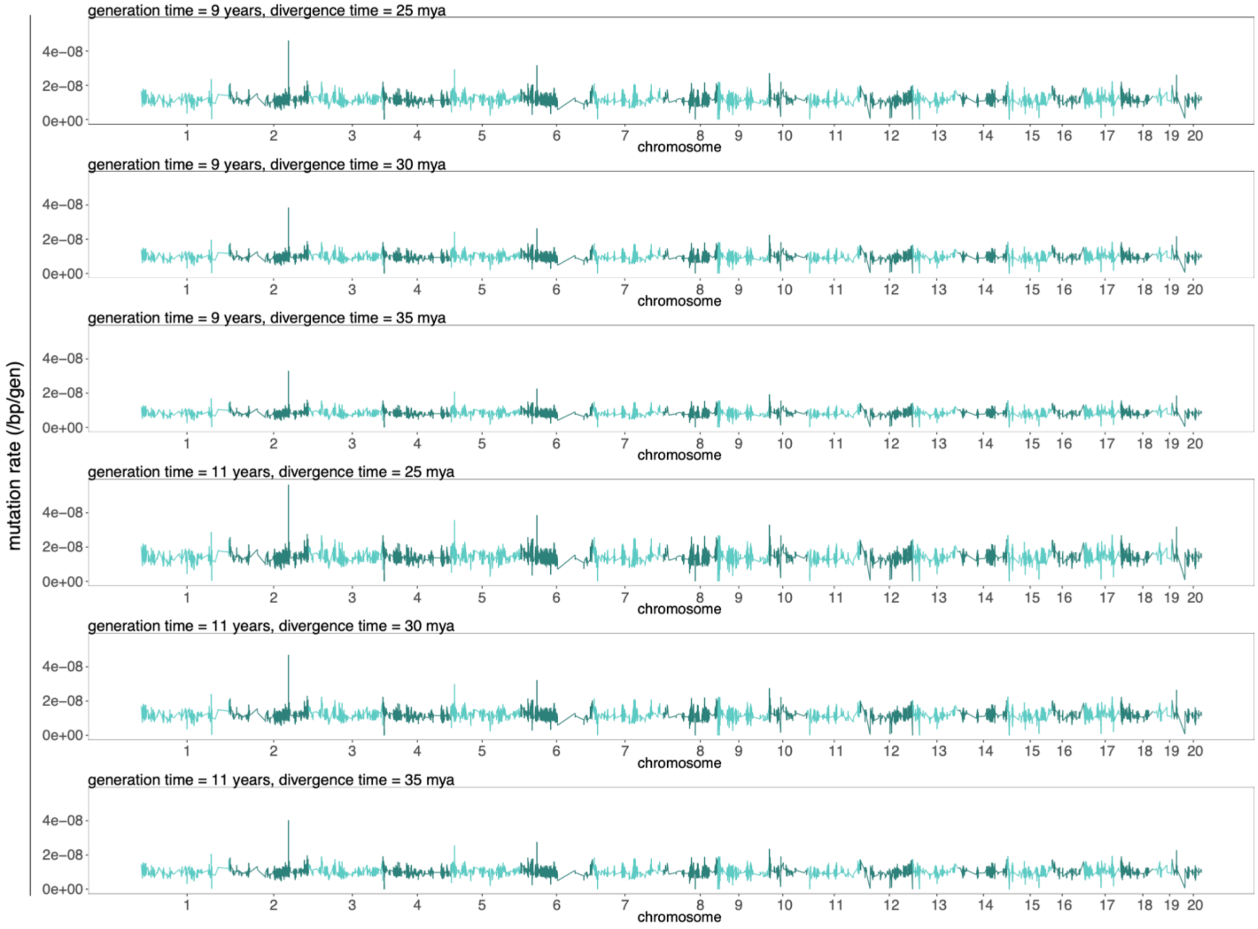


**Figure S3.** Fine-scale per-site per-generation mutation rate estimates observed in 10 kb windows with a step size of 5 kb based on patterns of neutral divergence between *M. mulatta* and *H. sapiens*, assuming generation times of 9 years and 11 years (Gagliardi et al. 2007; Xue et al. 2016) as well as divergence times ranging from 25 mya to 35 mya for *M. mulatta*–*H. sapiens* (Goodman et al. 1998; and see the reviews by Disotell and Tosi 2007; Skipper 2007; Chintalapati and Moorjani 2020). Individual chromosomes are shown in alternating shading.

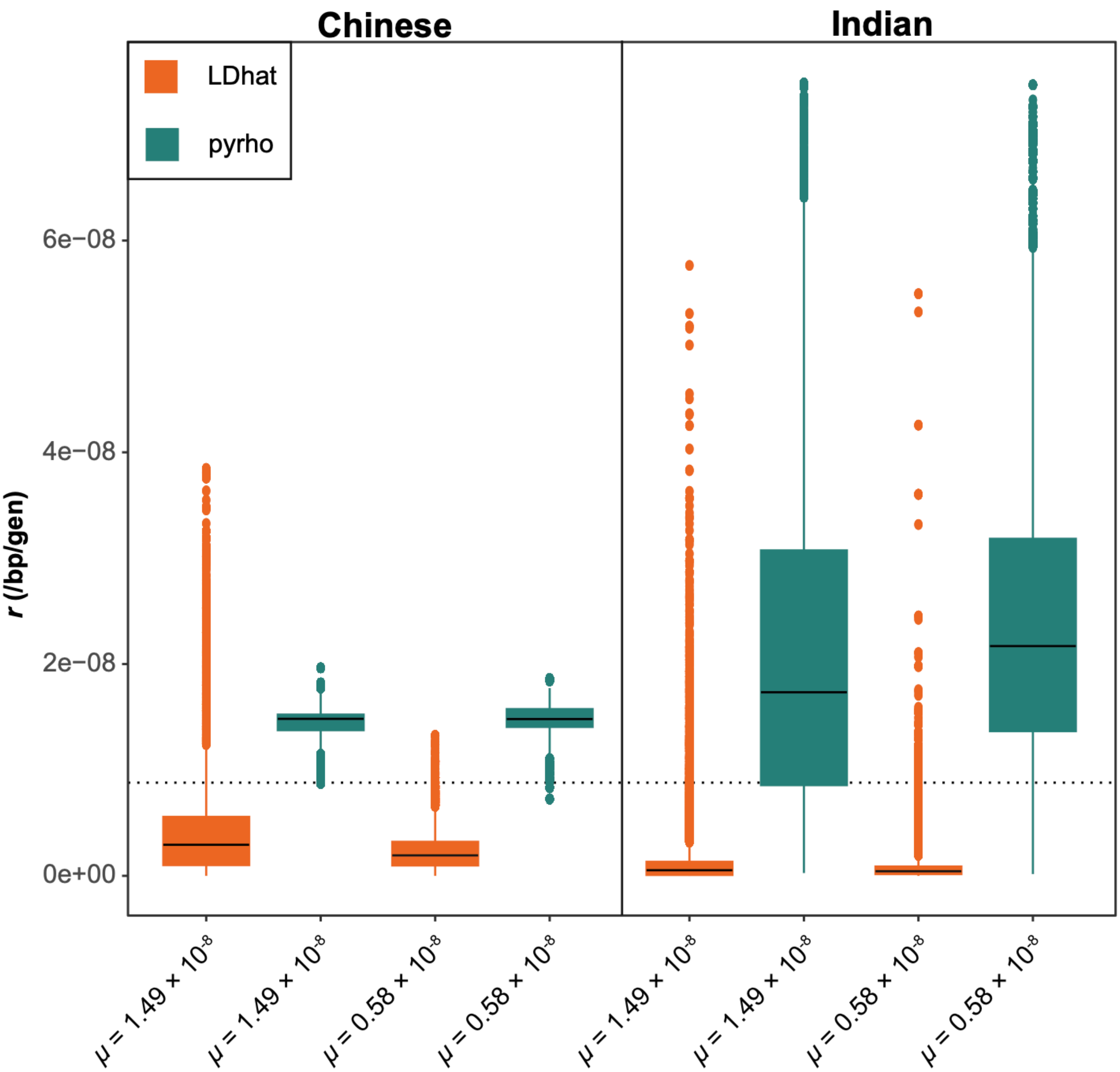


**Figure S4.** Comparison of per-site recombination rate estimates obtained from LDhat (shown in orange) and pyrho (teal) using simulated datasets of the Chinese and Indian population histories as inferred by Heenkenda et al. (2026), assuming a constant recombination rate of 0.88 cM/Mb (as inferred by Versoza et al. [2024] from pedigree data; indicated by a horizontal dashed line) as well as a constant mutation rate of either $0.58 \times 10^{-8}$ /bp/gen (i.e., the mean pedigree-based estimate of Wang et al. [2020]) or $1.49 \times 10^{-8}$ /bp/gen (i.e., the maximum indirect estimate observed from patterns of *M. mulatta–H. sapiens* divergence in this study).

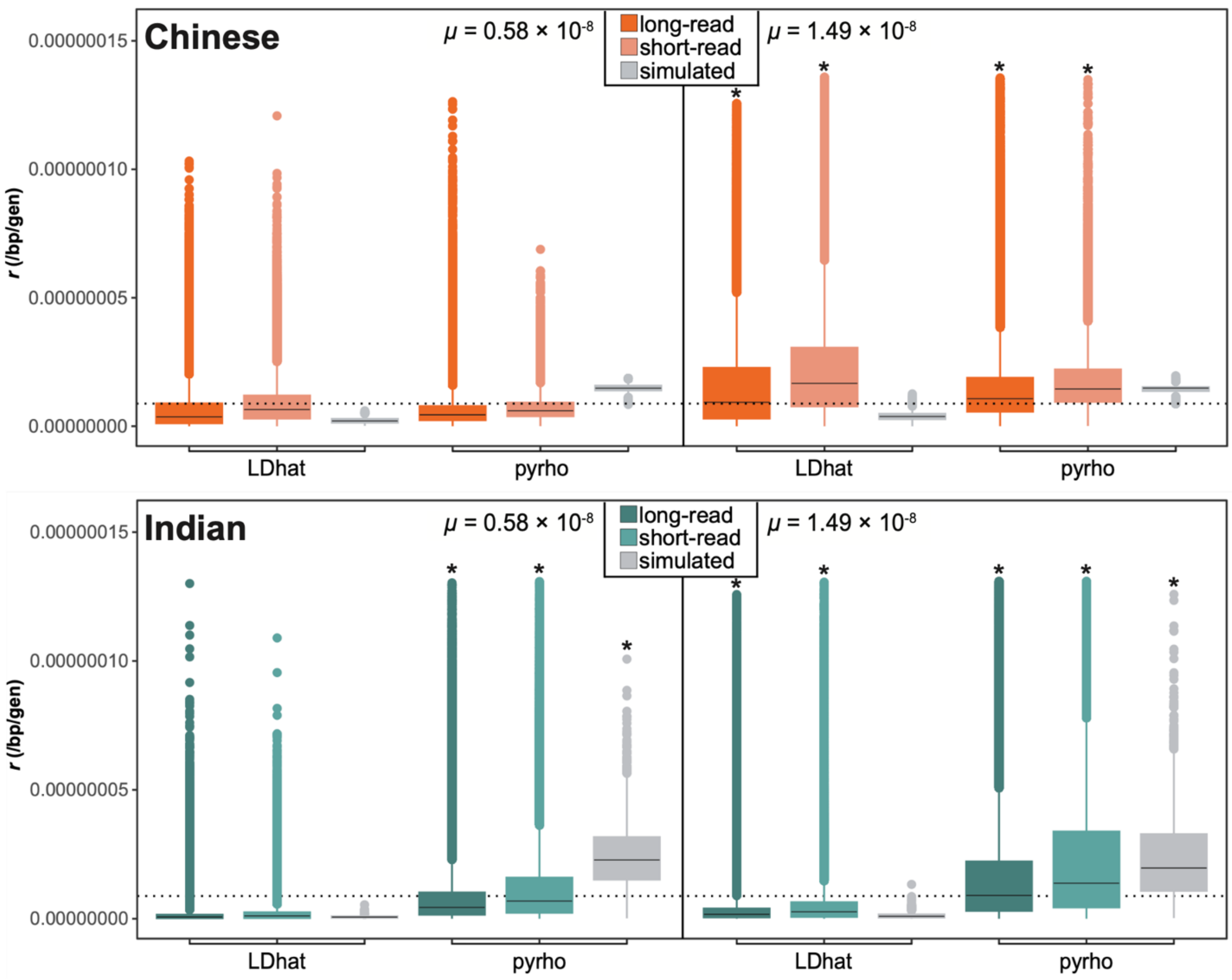


**Figure S5**. Per-site per-generation recombination rate estimates for rhesus macaque populations of (top) Chinese descent and (bottom) Indian descent as inferred using LDhat and pyrho at the 10kb-scale across the genome. Estimates obtained from empirical long-read and short-read data are shown dark and light colors, respectively (orange: Chinese; teal: Indian). For comparison, estimates obtained from simulations under the population-specific demographic histories recently inferred by Heenkenda et al. (2026) assuming a constant recombination rate (0.88 cM/Mb; as inferred by Versoza et al. [2024] from pedigrees; indicated by a dotted line) are shown in gray. Population-scaled recombination rate estimates were converted to per-generation estimates using population-specific $N_e$ values based on mutation rates ($\mu$) of $0.58 \times 10^{-8}$ /bp/gen (i.e., the pedigree-based estimate obtained by Wang et al. [2020]) or $1.49 \times 10^{-8}$ /bp/gen (i.e., the maximum indirect estimate observed from patterns of *M. mulatta*–*H. sapiens* divergence in this study). Asterisks indicate where observations were truncated for visualization.

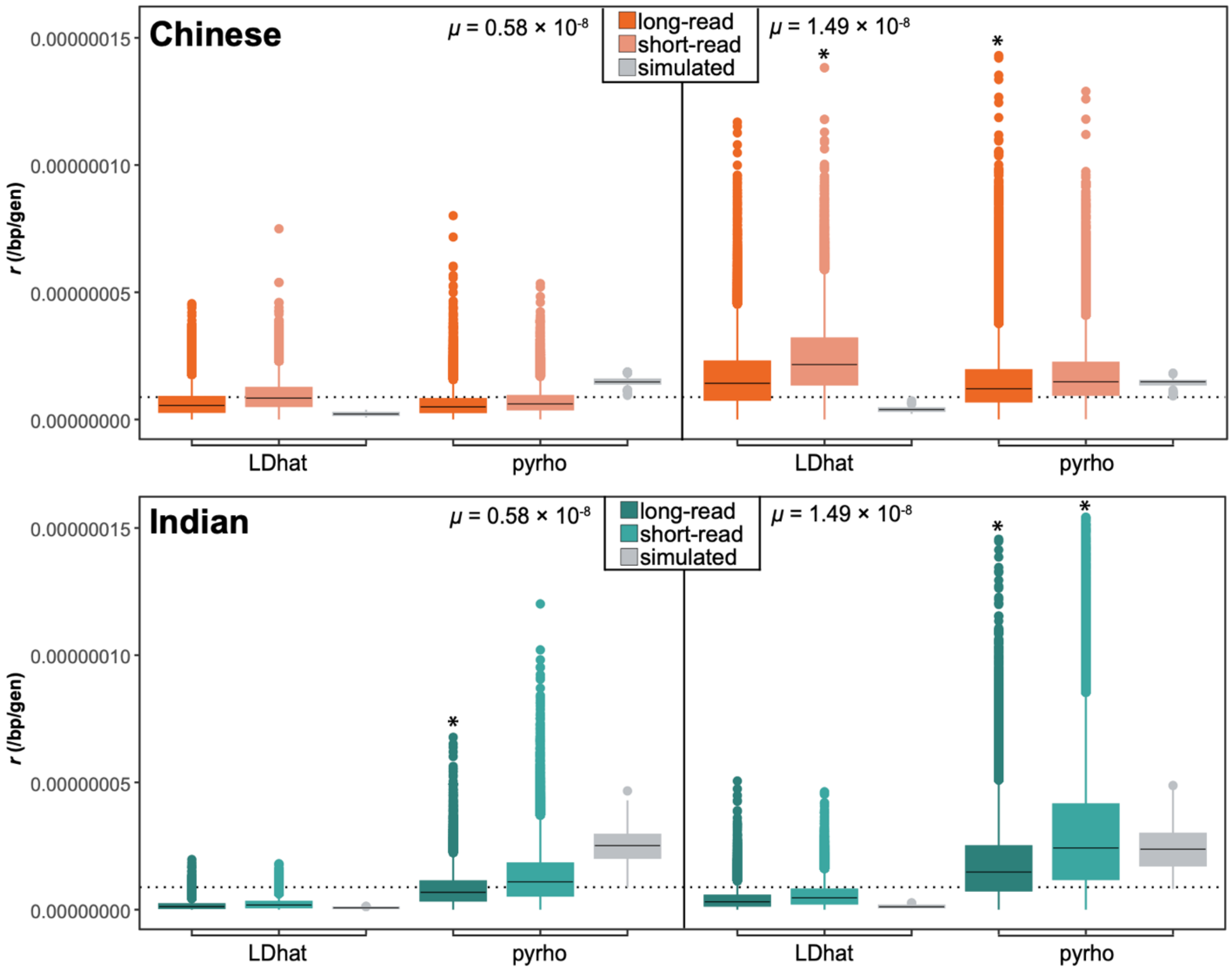


**Figure S6**. Per-site per-generation recombination rate estimates for rhesus macaque populations of (top) Chinese descent and (bottom) Indian descent as inferred using LDhat and pyrho at the 100kb-scale across the genome. Estimates obtained from empirical long-read and short-read data are shown dark and light colors, respectively (orange: Chinese; teal: Indian). For comparison, estimates obtained from simulations under the population-specific demographic histories recently inferred by Heenkenda et al. (2026) assuming a constant recombination rate (0.88 cM/Mb; as inferred by Versoza et al. [2024] from pedigrees; indicated by a dotted line) are shown in gray. Population-scaled recombination rate estimates were converted to per-generation estimates using population-specific $N_e$ values based on mutation rates ($\mu$) of $0.58 \times 10^{-8}$ /bp/gen (i.e., the pedigree-based estimate obtained by Wang et al. [2020]) or $1.49 \times 10^{-8}$ /bp/gen (i.e., the maximum indirect estimate observed from patterns of *M. mulatta*–*H. sapiens* divergence in this study). Asterisks indicate where observations were truncated for visualization.

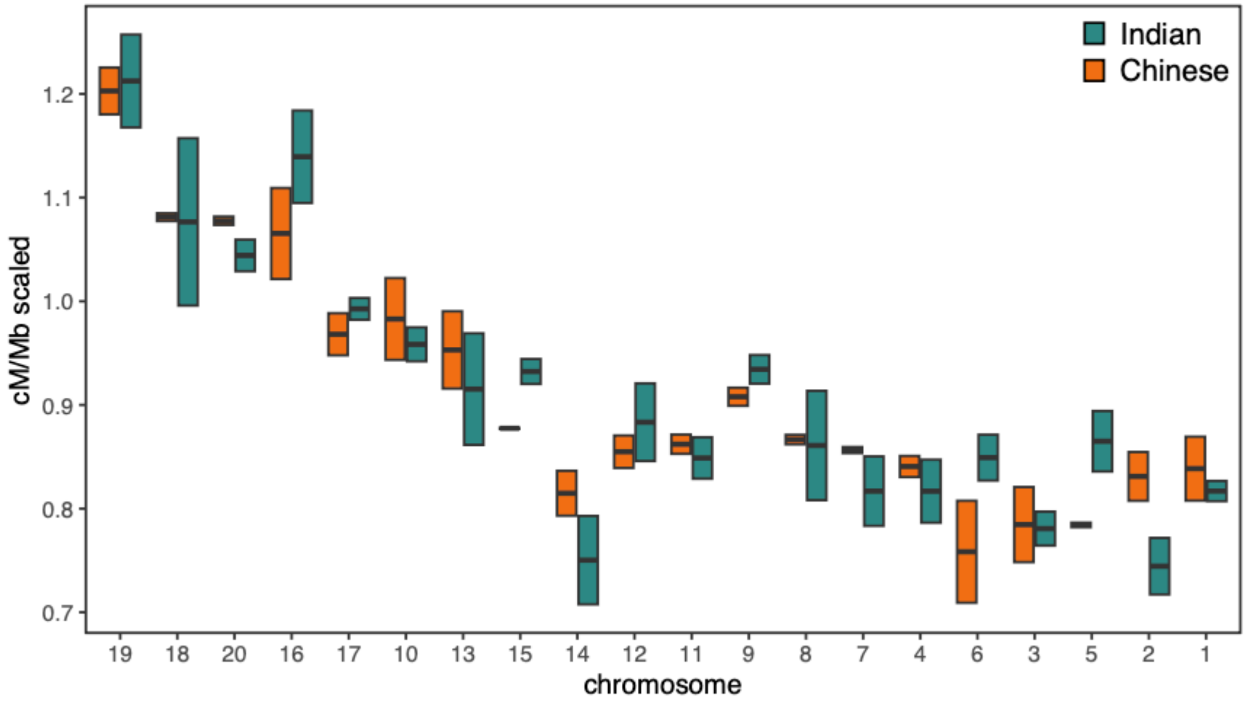


**Figure S7**. Per-chromosome recombination rate estimates for rhesus macaque populations of Chinese descent (shown in orange) and Indian descent (teal) as inferred using LDhat at the 1Mb-scale. Population-scaled recombination rate estimates were converted to per-generation estimates using population-specific $N_e$ values based on mutation rates ($\mu$) of $0.58 \times 10^{-8}$ /bp/gen (i.e., the pedigree-based estimate obtained by Wang et al. [2020]) or $1.49 \times 10^{-8}$ /bp/gen (i.e., the maximum indirect estimate observed from patterns of *M. mulatta*–*H. sapiens* divergence in this study) and re-scaled to the genetic map length previously observed from pedigree data (Versoza et al. 2024). Chromosomes are ordered from smallest to largest (left to right).

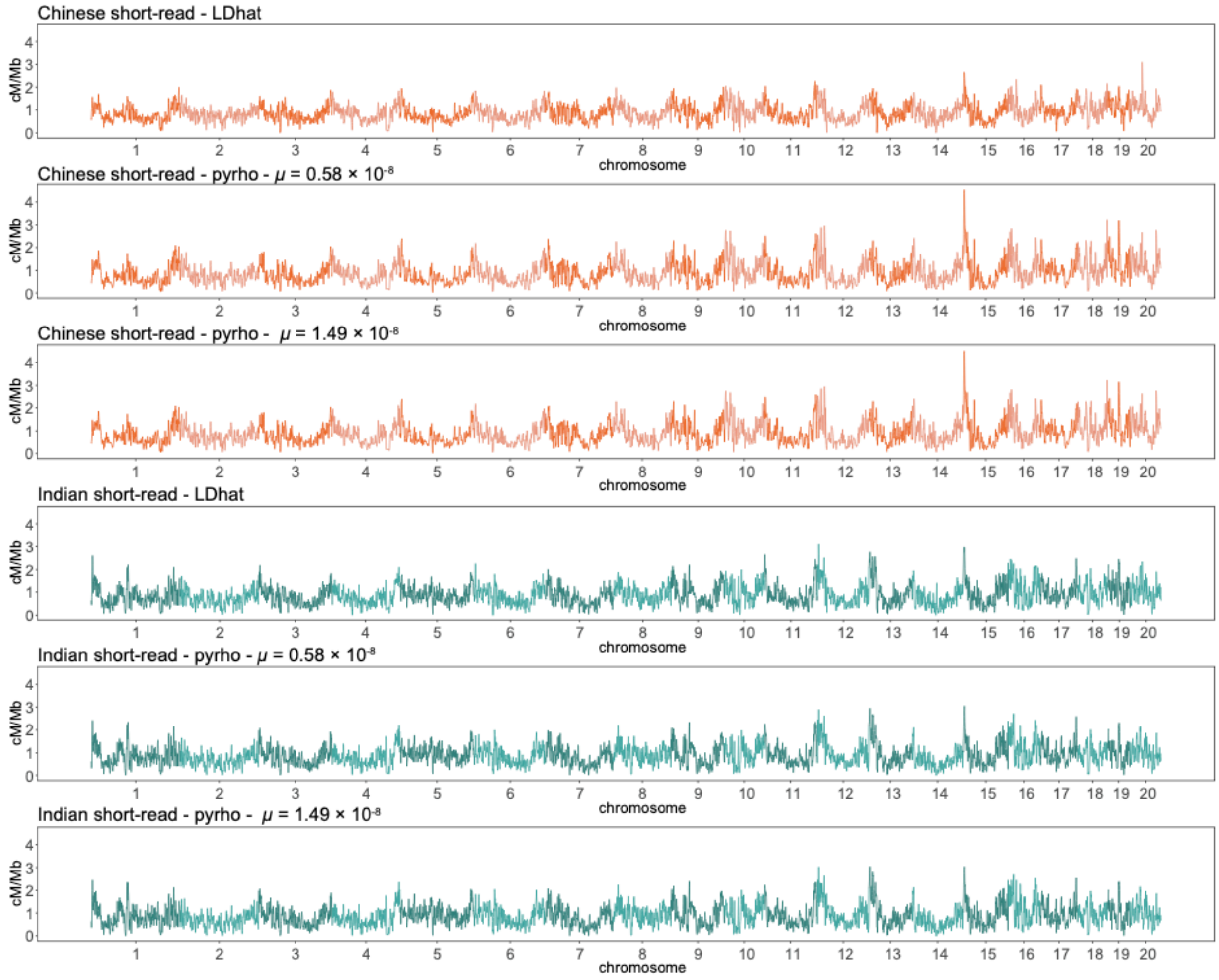


**Figure S8.** Fine-scale recombination rate maps for rhesus macaque populations of Chinese descent (shown in orange) and Indian descent (teal) as inferred from short-read data using LDhat and pyrho at the 1Mb-scale across the genome. Population-scaled recombination rate estimates were converted to per-generation estimates using population-specific $N_e$ values based on mutation rates ($\mu$) of $0.58 \times 10^{-8}$ /bp/gen (i.e., the pedigree-based estimate obtained by Wang et al. [2020]) or $1.49 \times 10^{-8}$ /bp/gen (i.e., the maximum indirect estimate observed from patterns of *M. mulatta–H. sapiens* divergence in this study) and re-scaled to the genetic map length previously observed from pedigree data (Versoza et al. 2024). Individual chromosomes are shown in alternating shading.

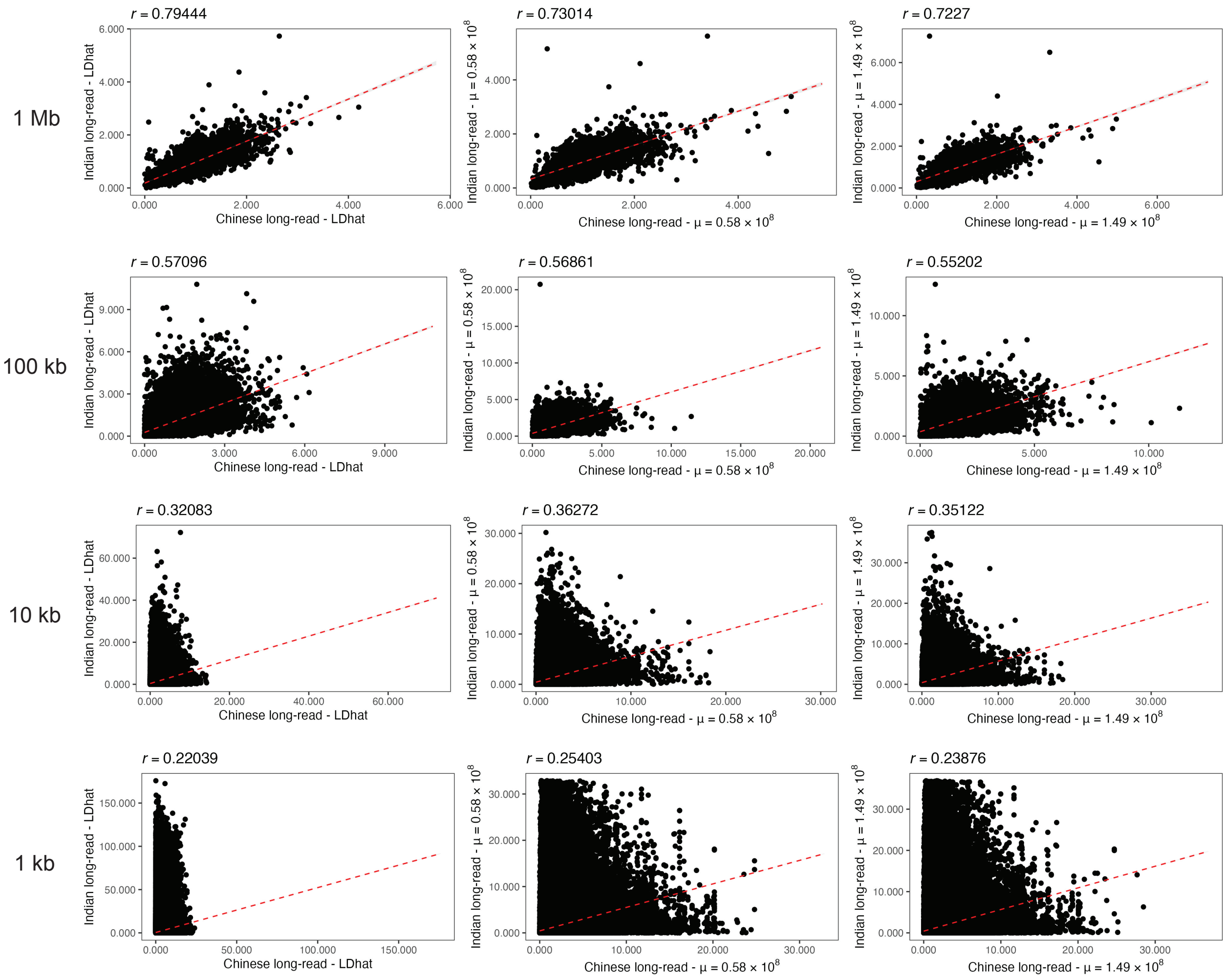


**Figure S9.** Pearson correlation coefficients (*r*) between recombination rate maps for Chinese and Indian populations inferred from long-read data using LDhat (left panels) and pyrho (middle and right panels) at 1Mb-, 100kb-, 10kb-, and 1kb-scales. Population-scaled recombination rate estimates were converted to per-generation estimates using population-specific $N_e$ values based on mutation rates ($\mu$) of 0.58 $\times$ $10^{-8}$ /bp/gen (i.e., the pedigree-based estimate obtained by Wang et al. [2020]) or 1.49 $\times$ $10^{-8}$ /bp/gen (i.e., the maximum indirect estimate observed from patterns of *M. mulatta*–*H. sapiens* divergence in this study) and re-scaled to the genetic map length previously observed from pedigree data (Versoza et al. 2024).

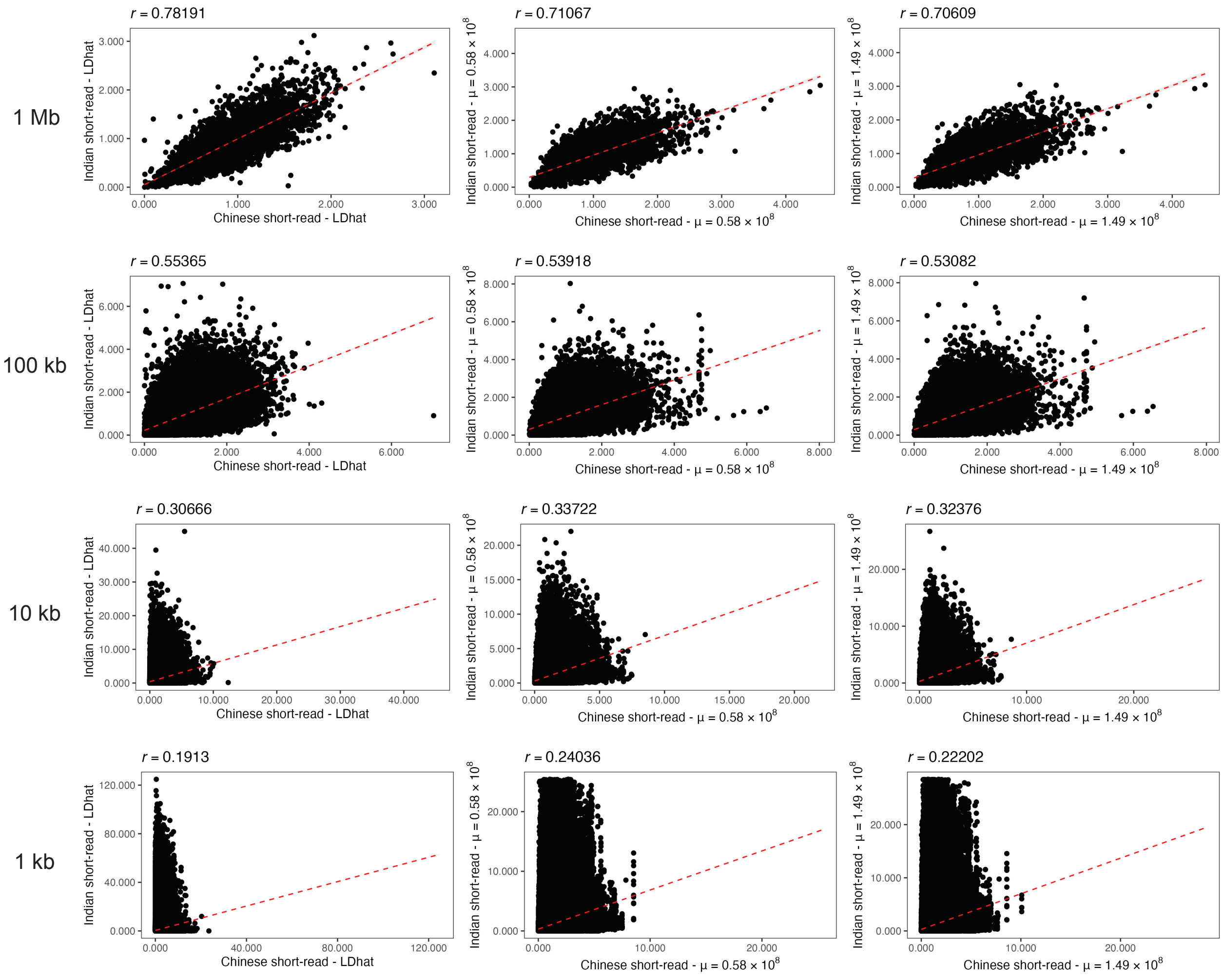


**Figure S10.** Pearson correlation coefficients ($r$) between recombination rate maps for Chinese and Indian populations inferred from short-read data using LDhat (left panels) and pyrho (middle and right panels) at 1Mb-, 100kb-, 10kb-, and 1kb-scales. Population-scaled recombination rate estimates were converted to per-generation estimates using population-specific $N_e$ values based on mutation rates ($\mu$) of 0.58 $\times$ $10^{-8}$ /bp/gen (i.e., the pedigree-based estimate obtained by Wang et al. [2020]) or 1.49 $\times$ $10^{-8}$ /bp/gen (i.e., the maximum indirect estimate observed from patterns of *M. mulatta*–*H. sapiens* divergence in this study) and re-scaled to the genetic map length previously observed from pedigree data (Versoza et al. 2024).

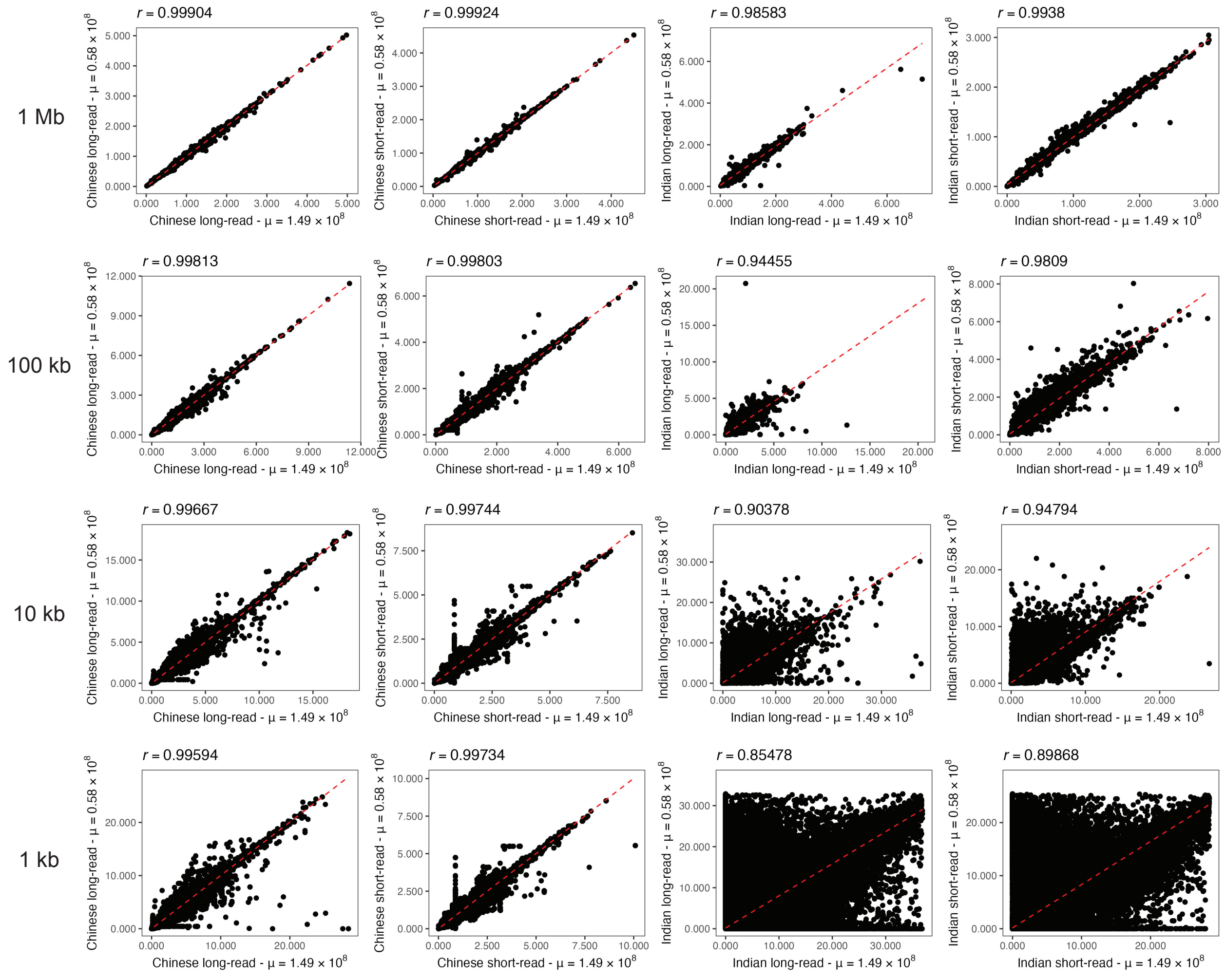


**Figure S11.** Pearson correlation coefficients (*r*) between recombination rate maps inferred in pyrho from the same read type (i.e., long- or short-reads) but under different demographic models for the rhesus macaque populations of Chinese descent (two left-most panels) and Indian descent (two right-most panels) at 1Mb-, 100kb-, 10kb-, and 1kb-scales. Population-scaled recombination rate estimates were converted to per-generation estimates using population-specific $N_e$ values based on mutation rates ($\mu$) of $0.58 \times 10^{-8}$ /bp/gen (i.e., the pedigree-based estimate obtained by Wang et al. [2020]) or $1.49 \times 10^{-8}$ /bp/gen (i.e., the maximum indirect estimate observed from patterns of *M. mulatta–H. sapiens* divergence in this study) and re-scaled to the genetic map length previously observed from pedigree data (Versoza et al. 2024).

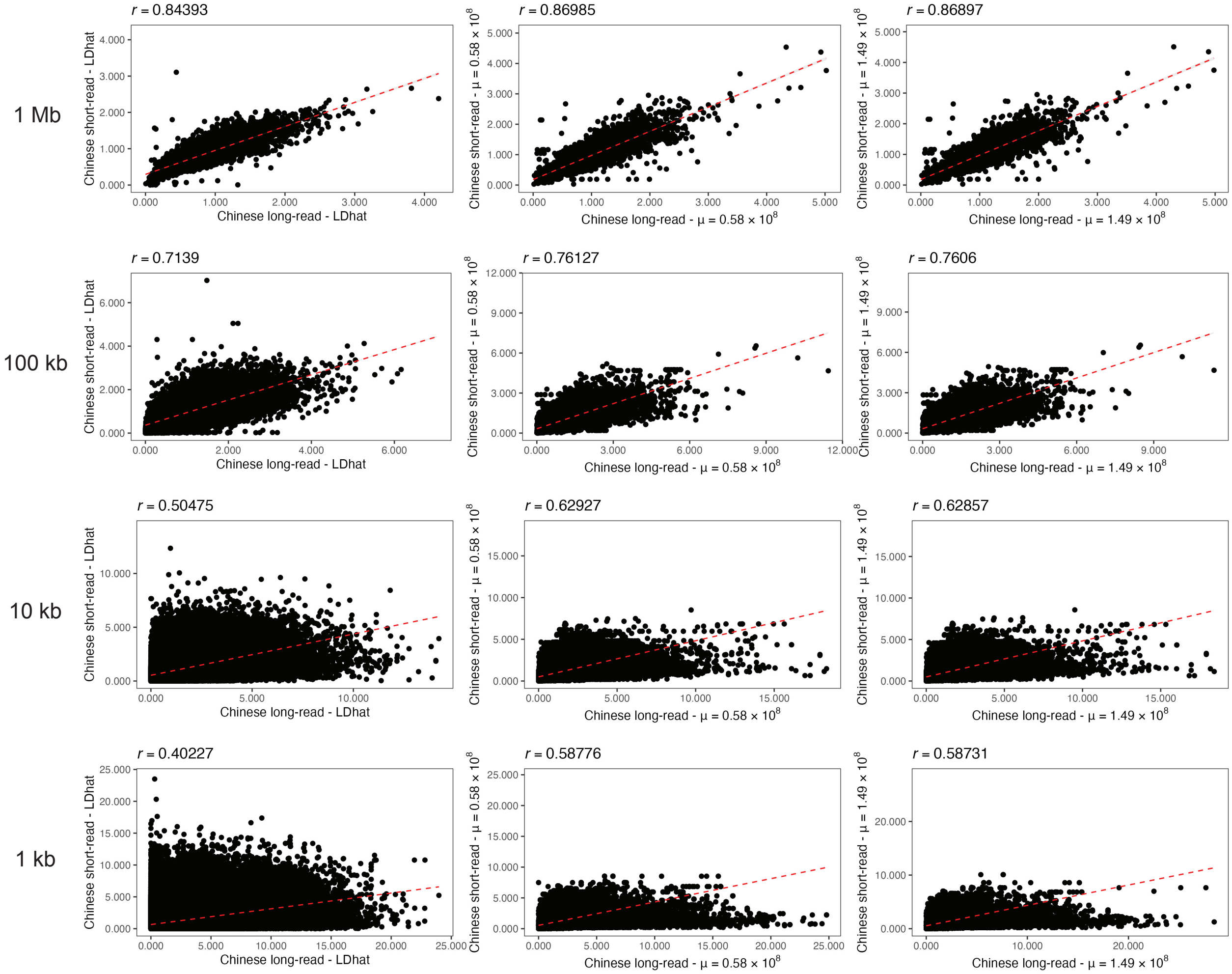


**Figure S12.** Pearson correlation coefficients (*r*) between recombination rate maps inferred from long-read and short-read data using LDhat (left panels) and pyrho (middle and right panels) for the Chinese population at 1Mb-, 100kb-, 10kb-, and 1kb-scales. Population-scaled recombination rate estimates were converted to per-generation estimates using population-specific $N_e$ values based on mutation rates ($\mu$) of $0.58 \times 10^{-8}$ /bp/gen (i.e., the pedigree-based estimate obtained by Wang et al. [2020]) or $1.49 \times 10^{-8}$ /bp/gen (i.e., the maximum indirect estimate observed from patterns of *M. mulatta*–*H. sapiens* divergence in this study) and re-scaled to the genetic map length previously observed from pedigree data (Versoza et al. 2024).

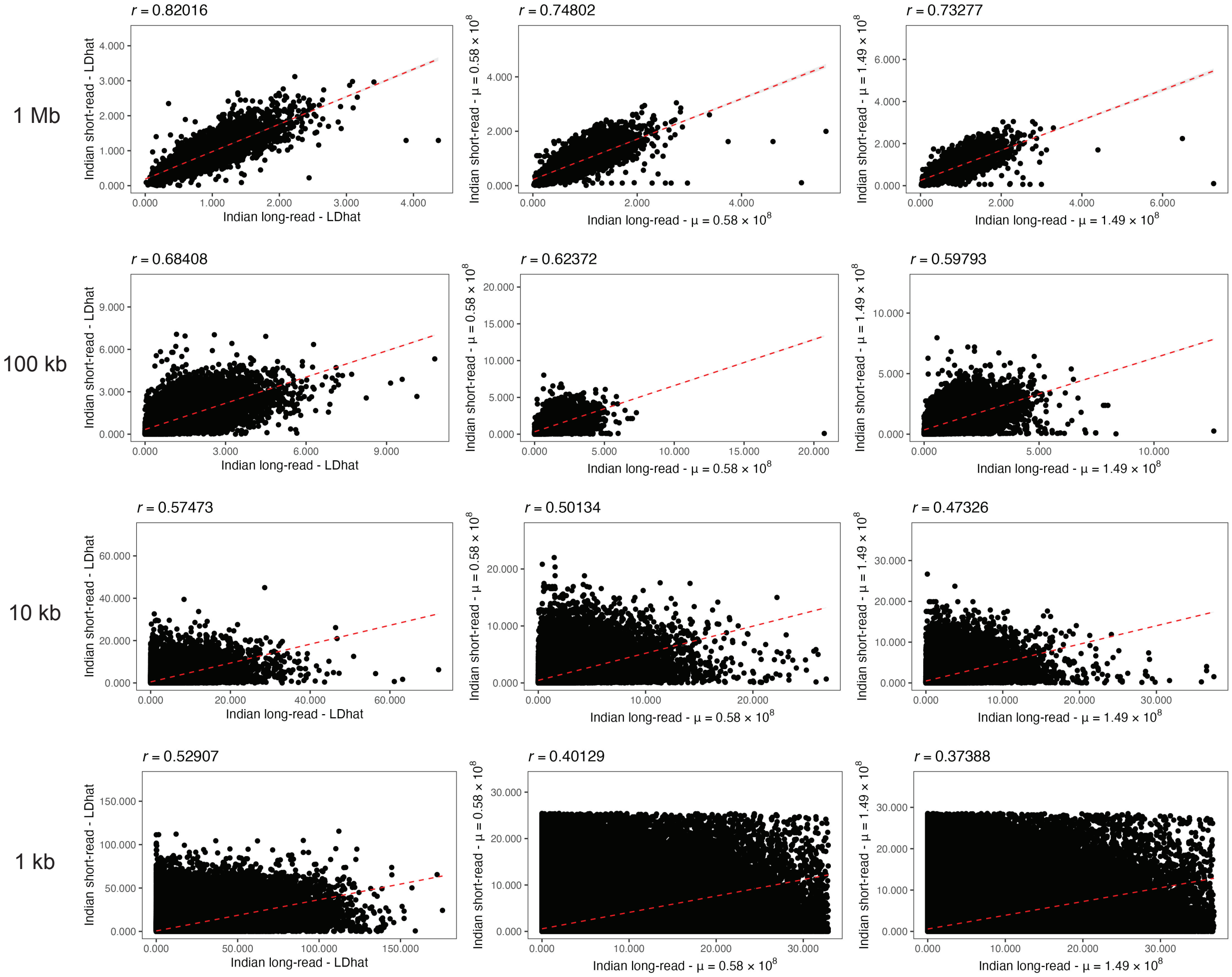


**Figure S13.** Pearson correlation coefficients (*r*) between recombination rate maps inferred from long-read and short-read data using LDhat (left panels) and pyrho (middle and right panels) for the Indian population at 1Mb-, 100kb-, 10kb-, and 1kb-scales. Population-scaled recombination rate estimates were converted to per-generation estimates using population-specific $N_e$ values based on mutation rates ($\mu$) of 0.58 × $10^{-8}$ /bp/gen (i.e., the pedigree-based estimate obtained by Wang et al. [2020]) or 1.49 × $10^{-8}$ /bp/gen (i.e., the maximum indirect estimate observed from patterns of *M. mulatta*–*H. sapiens* divergence in this study) and re-scaled to the genetic map length previously observed from pedigree data (Versoza et al. 2024).

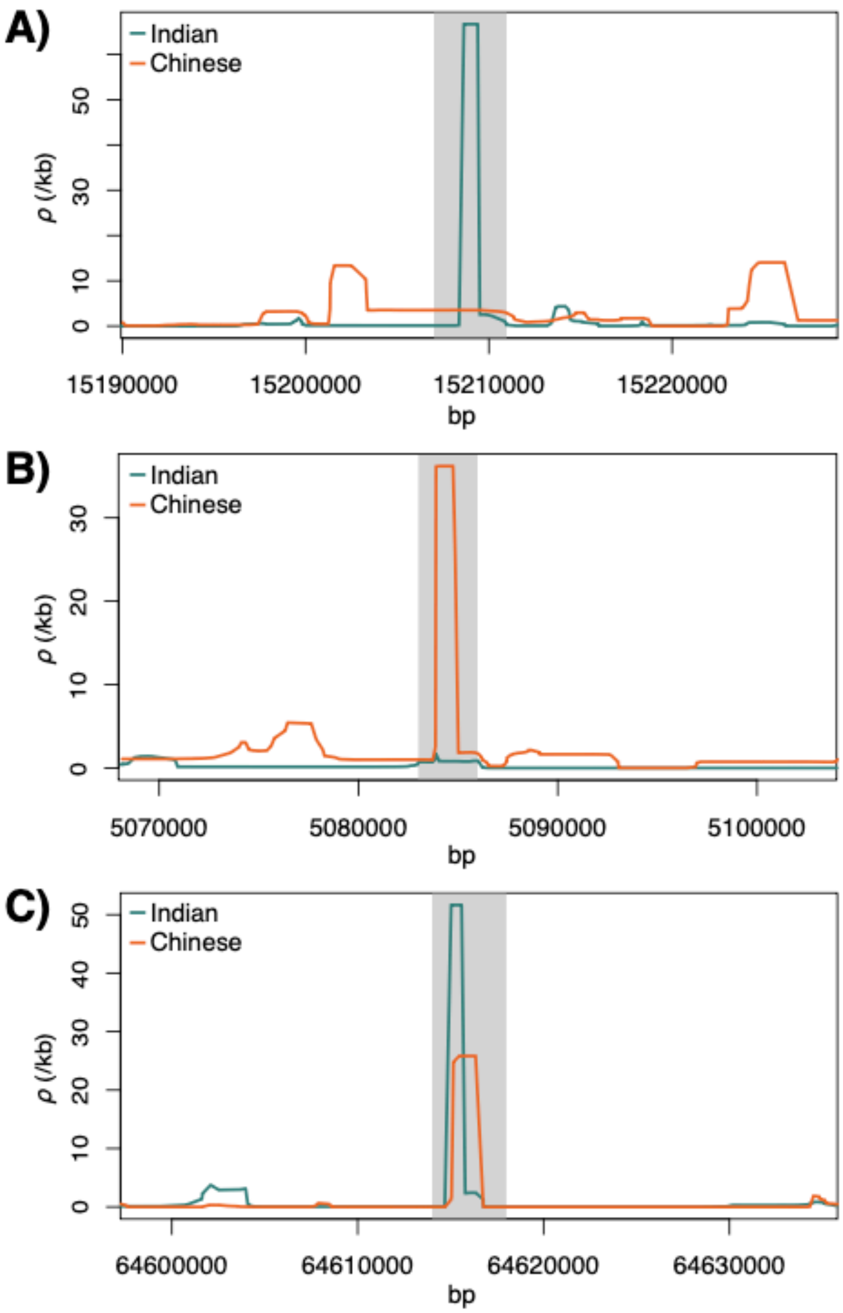


**Figure S14**. Examples of recombination hotspots **(A)** unique to the Indian rhesus macaque population, **(B)** unique to the Chinese rhesus macaque population, and **(C)** shared between both populations.
